# Hidden route of protein damage through confined oxygen gas

**DOI:** 10.1101/2024.01.03.574110

**Authors:** Seoyoon Kim, Eojin Kim, Mingyu Park, Seong Ho Kim, Byung-Gyu Kim, Victor W. Sadongo, W.C. Bhashini Wijesinghe, Yu-Gon Eom, Gwangsu Yoon, Chaiheon Lee, Hannah Jeong, Chae Un Kim, Kyungjae Myung, Jeong-Mo Choi, Seung Kyu Min, Tae-Hyuk Kwon, Duyoung Min

**Affiliations:** Department of Chemistry, Ulsan National Institute of Science and Technology, Ulsan 44919, Republic of Korea; Center for Genomic Integrity, Institute for Basic Science, Ulsan 44919, Republic of Korea; Department of Chemistry and Chemistry Institute for Functional Materials, Pusan National University, Busan 46241, Republic of Korea; Department of Physics, Ulsan National Institute of Science and Technology, Ulsan 44919, Republic of Korea; Department of Biomedical Engineering, Ulsan National Institute of Science and Technology, Ulsan 44919, Republic of Korea; Center for Wave Energy Materials, Ulsan National Institute of Science and Technology, Ulsan 44919, Republic of Korea

**Keywords:** protein oxidative damage, alternative photooxidation pathway, protein cavities, oxygen confinement, singlet oxygen generation, protein core oxidation

## Abstract

Oxidative modifications can severely impair protein structure, fold, and function, closely linked to human aging and diseases. Conventional oxidation pathways typically involve the free diffusion of reactive oxygen species (ROS), followed by chemical attacks on the protein surface. Here, we report a hidden route of protein oxidative damage, which we refer to as O_2_-confinement oxidation pathway. This pathway starts with the initial trapping of dissolved molecular oxygen (O_2_) within protein cavity spaces, followed by interaction with photosensitizing tryptophan residues. The trapped O_2_ is then converted to singlet oxygen (^1^O_2_), a powerful ROS, through spin-flip electron transfer mechanism under blue light. The generated ^1^O_2_ within the protein ultimately attacks the protein core residues through constrained diffusion, accelerating the oxidative damage. This alternative photooxidation pathway through the initial O_2_ trapping would bypass the antioxidant defense systems which target freely-diffusing ROS, constituting an additional layer of protein oxidative damage in cells and tissues.

---

Protein integrity in living organisms can be compromised by nonenzymatic posttranslational modifications under both physiological and pathological conditions^1–4^. Protein quality control systems, mainly involving proteasomal and lysosomal pathways, eliminate proteins with impaired folds and functions^5–9^. However, oxidative stress caused by redox imbalance leads to the accumulation of defective proteins and cellular dysfunction, associated with human aging and diseases such as neurodegenerative disorders and cancers^7,10–19^.

The primary pathways of protein oxidation involve the conversion to reactive oxygen species (ROS), followed by their free diffusion in cells or solutions, and subsequent oxidation of easily accessible protein surface residues^11,16,20–22^. Endogenous ROS is typically generated as natural byproducts of cellular metabolism, such as aerobic respiration^14,23–25^. Exposure of light, particularly within the ultraviolet-visible light spectrum, can also lead to ROS generation through energy transfer or electron transfer mediated by photosensitizing molecules^26,27^.

Of special note is that molecular oxygen (O_2_) can dissolve in human tissues up to a considerable level of ∼13% as pO_2_ (ref. ^28^) and can directly bind to local protein structures, even in the absence of transition metal ions^29,30^. However, it is entirely unknown whether and how the direct O_2_ binding to proteins leads to their oxidative damage. This oxidation pathway could be a crucial factor to consider in maintaining protein stability and function, designing more robust proteins resistant to oxidation, and understanding the aging and pathophysiology associated with protein oxidation.

In this study, we elucidate a unique O_2_-confinement pathway/mechanism of protein photooxidation. Using a single-molecule tweezer approach under blue light, we found that contrary to expectations, protein oxidative damage was facilitated even in its folded state that conceals a substantial portion of oxidizable atoms. Various methods – mass spectrometry analysis, spectrophotometric assays, electron paramagnetic resonance spectroscopy, molecular dynamics simulations, first-principles calculations, and Monte Carlo analysis – collectively point to the O_2_-confinement oxidation we suggest here as the primary cause of the unusual oxidative damage. In this alternative photooxidation pathway, singlet oxygen (^1^O_2_,), a potent ROS, is generated within the folded structure and subsequently attacks the protein core residues through constrained diffusion (Fig. 1). The ^1^O_2_ generation is mediated by the blue-light excitation of the molecular system comprising the trapped O_2_ and nearby tryptophan (Trp) residue. Our study suggests that the internal architecture of protein folds partially encodes the inherent susceptibility to oxidative damage.

**Fig. 1.**
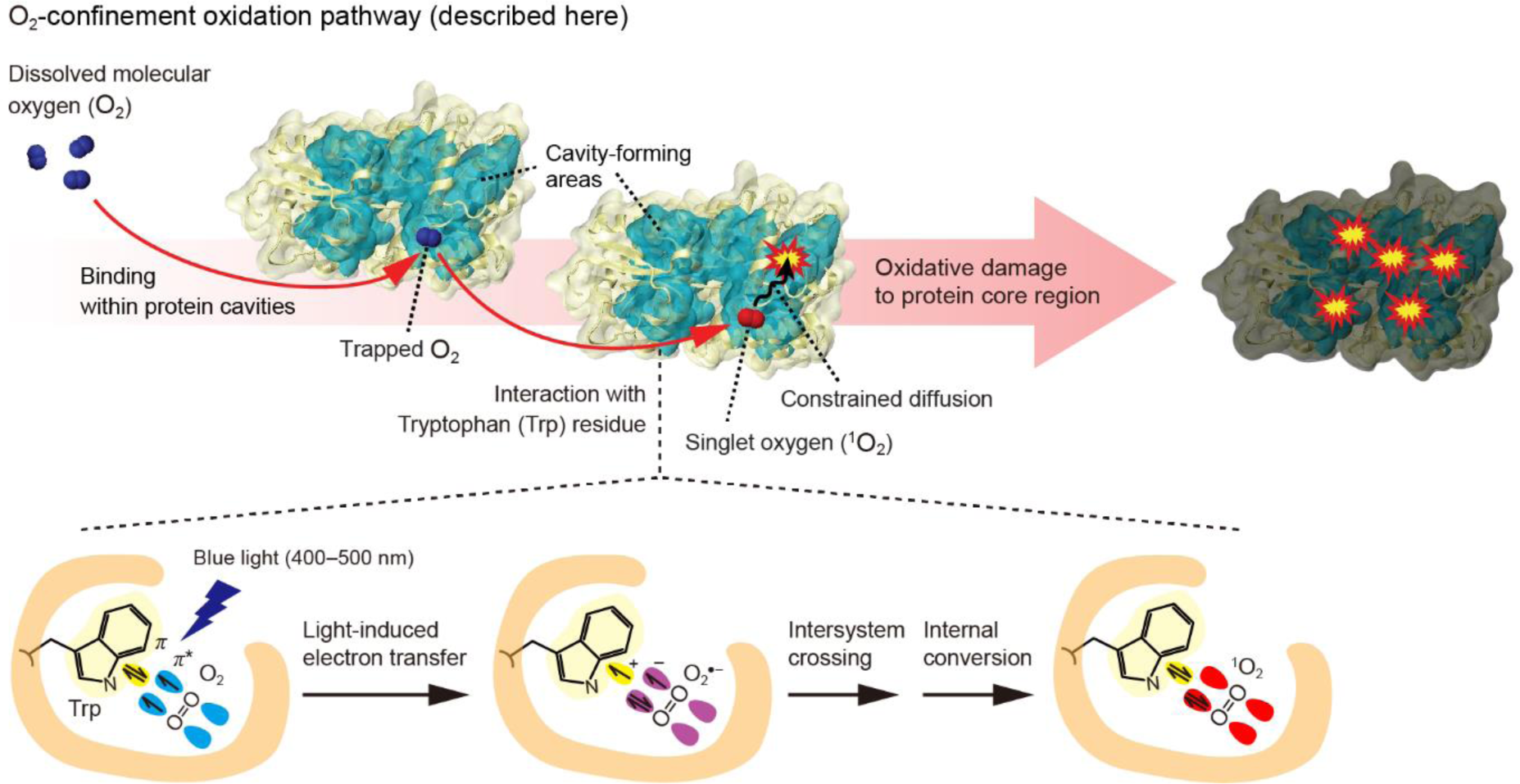
| Proposed model of protein oxidative damage through O_2_-confinement pathway. Dissolved oxygen gas (O_2_) is trapped within protein cavities containing tryptophan (Trp) as an endogenous photosensitizer. Upon exposure to blue light (400–500 nm), the trapped O_2_ molecule undergoes conversion to singlet oxygen (^1^O_2_), a potent reactive oxygen species (ROS), through spin-flip electron transfer mechanism. The generated ^1^O_2_ oxidizes nearby residues through constrained diffusion within the protein.

## Results

### Declining protein foldability during cyclic unfolding

This study was prompted by the unexpected observation of a rapid decline in protein foldability under blue light (λ_peak_ = 447 nm; Extended Data Fig. 1 for the full spectrum), using a recently developed single-molecule tweezers^31^ (Fig. 2a). In this method, we tethered the N- and C-termini of a single protein to the glass surface and a magnetic bead using 1024-bp DNA handles through three orthogonal conjugation approaches: dibenzocyclooctyne-azide, traptavidin-2×biotin, and SpyCatcher-SpyTag (Methods and Extended Data Fig. 2a–c). This tweezer approach demonstrated high stability, enabling the capture of ∼1000 reversible unfolding events in a single protein^31^. We employed this method to repetitively unfold and refold maltose binding protein (MBP), a widely-used model protein in protein folding studies, over hundreds of cycles.

**Fig. 2.**
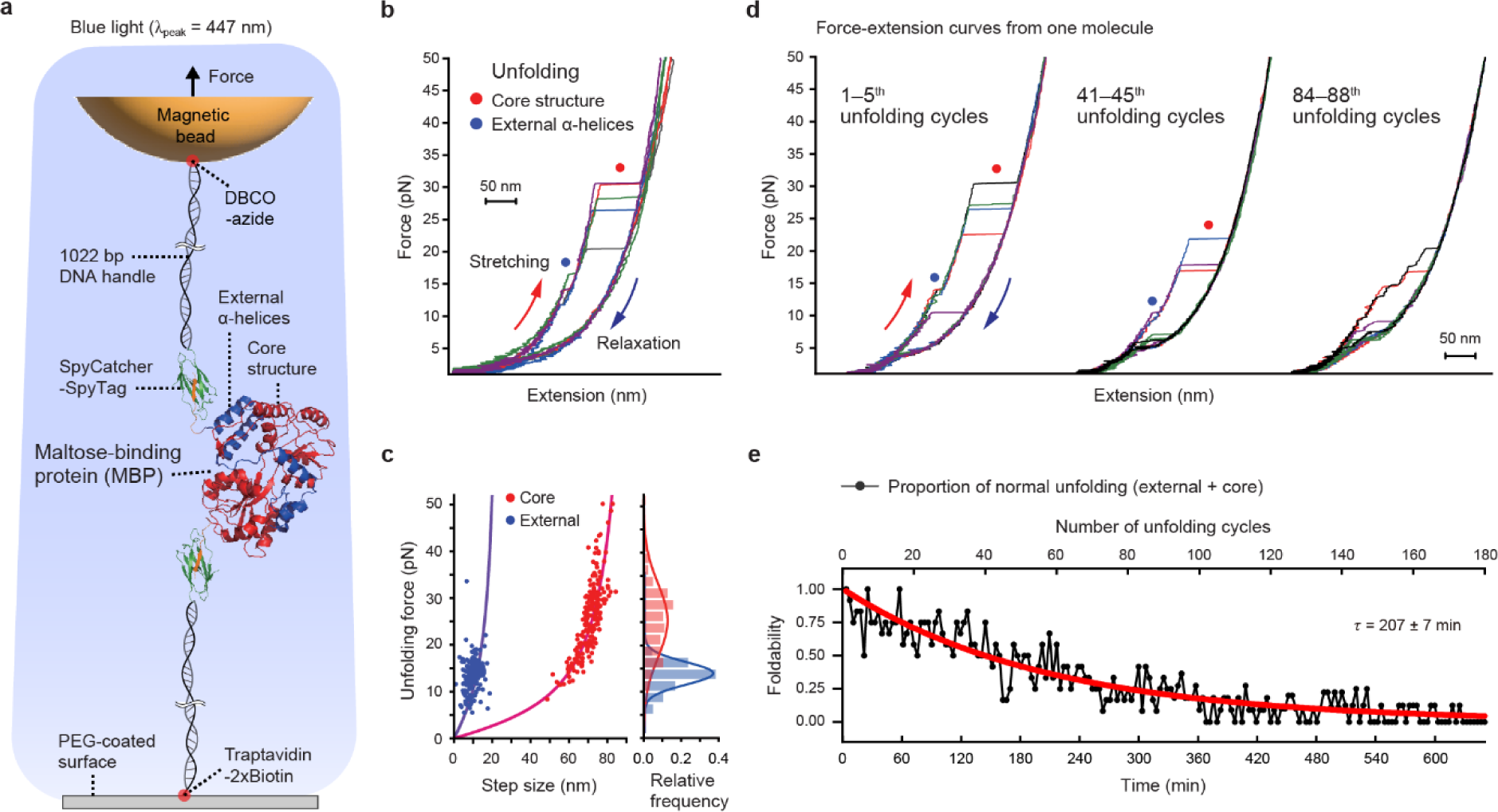
| Declining foldability of MBP during cyclic unfolding. (a) Schematics of our single-molecule magnetic tweezers. Maltose-binding protein (MBP) is repetitively unfolded over hundreds of cycles under blue light. (b) Representative force-extension curves of the protein-DNA hybrid molecule (*N* = 5 molecules). The blue and red dots point to the unfolding of the external α-helices at the C-terminus and the remaining core structure of MBP, respectively. (c) Scatter plot of unfolding forces and step sizes of MBP (*N* = 12 molecules, *n* = 203 data points for each structural region). The data were fitted by the worm-like chain model. The right histogram represents the relative frequency of unfolding forces. (d) Representative force-extension curves at different numbers of unfolding cycles (*n* = 5 for each cycle region from one molecule). (e) Foldability trend during cyclic unfolding. The foldability is quantified as the proportion of normal unfolding with the characteristic pattern, *i.e.*, the unfolding of the external α-helices followed by the core structure (*N* = 12 molecules). The corresponding times are also labeled on the bottom axis. The error of the decay time constant indicates SE.

During each unfolding cycle, the force was applied in the range of 1 pN to 50 pN and then immediately relaxed back to 1 pN, followed by a 2.5-min waiting to allow for complete refolding (Fig. 2b). In the initial several cycles, we clearly observed two distinct unfolding steps: a partial unfolding at lower forces followed by the complete unfolding of the remaining structure at higher forces (Fig. 2b). The distribution of forces and step sizes for each unfolding event was analyzed using worm-like chain model^32^ (Fig. 2c). The data were consistent with previous studies^33–36^, where the partial and complete unfolding corresponded to the external α-helices at the C-terminus and more stable core structure, respectively (Fig. 2a and Extended Data Fig. 2d).

As the number of unfolding cycles increased, however, the unfolding pattern became progressively inconsistent and difficult to define (Fig. 2d). This observation indicates the decrease in foldability to the native state. To quantitatively assess the declining foldability during the cyclic unfolding, we analyzed the proportion of normal unfolding (*i.e.*, the unfolding of the external α-helices followed by the core structure) at each number of unfolding cycles, obtained from multiple molecules (Fig. 2e). The protein foldability was found to decline to zero during approximately two hundred cycles, corresponding to a decay with a time constant (*τ*) of 207 ± 7 min. This result implies a gradual increase of defects in the primary structure, which serves as the starting point of folding into the tertiary structure in our experiments.

### Photooxidative damage as the cause of foldability decay

Throughout all the cycles, the refolding is driven by relaxing the stretched state of the identical fully unstructured polypeptide at 50 pN. Given this same initial state, the question arises about what factors contribute to the observed foldability decay. To elucidate the underlying causes, we first examined the potential influence of the repetitive pulling process itself (Fig. 3a, left). We compared the trends in the foldability upon a change in only one of the two pulling parameters: the pulling rate modulated by the magnet speed (0.3 *vs.* 0.01 mm/s) and the waiting time between pulling cycles (2.5 *vs.* 90 min). For all conditions, we found that the decay rates of foldability were comparable with *τ* = 207–226 min (Fig. 3b). This result indicates that the declining foldability during the cyclic unfolding is irrelevant to the repetitive process of pulling and unfolding.

**Fig. 3.**
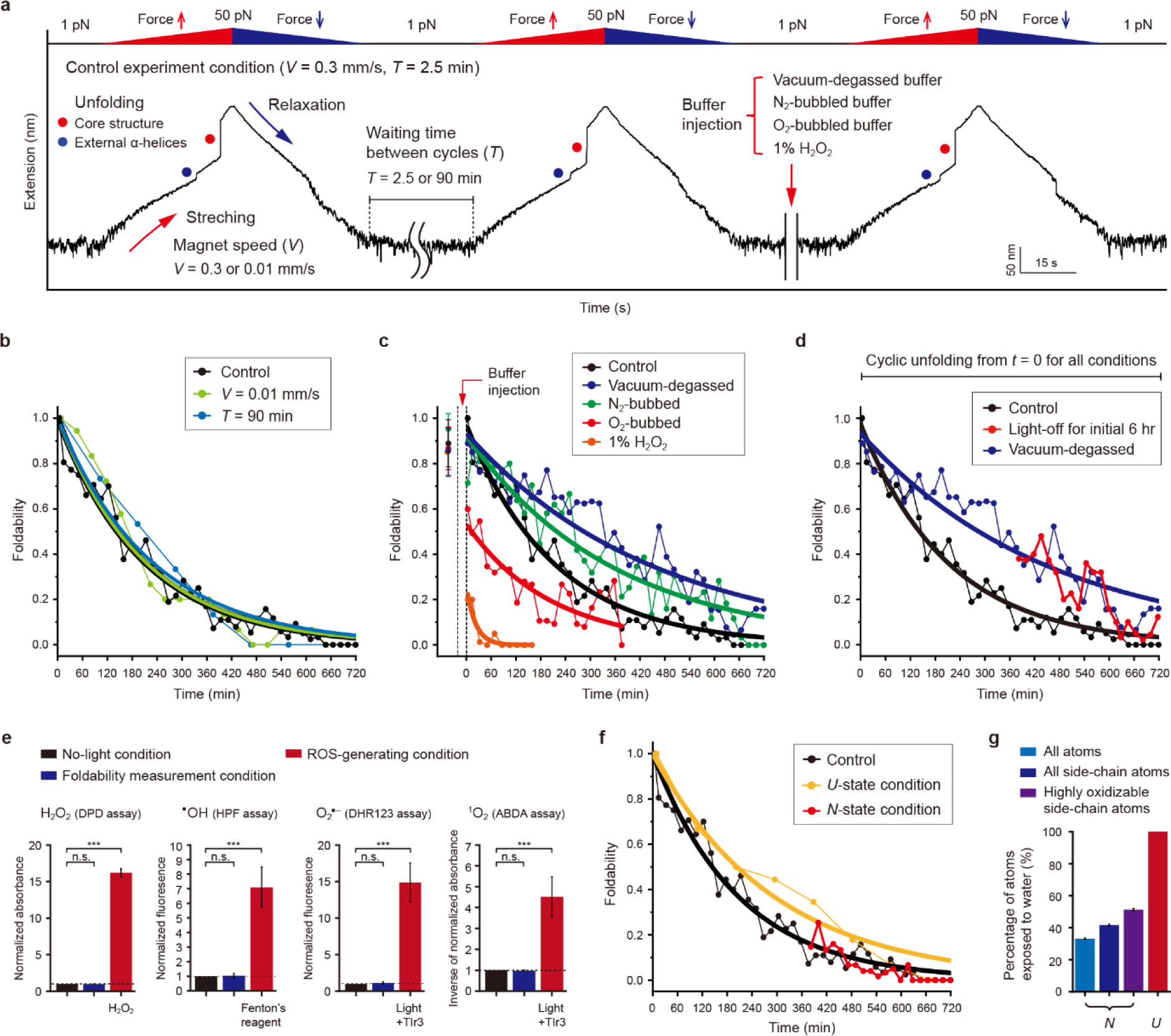
| Unusual photooxidative damage as main cause of the foldability decay. (a) Representative time-resolved extension trace showing various experimental conditions (*N* = 5– 12 molecules for each condition). (b–c) Foldability trend over time under different pulling conditions (b) or different oxidative stress conditions (c). In the latter experiments, each buffer is injected into the sample chamber after three unfolding cycles. (d) Foldability trend in the absence of blue light during the initial 6 hr. The cyclic unfolding is also carried out in the light-off condition during the entire time scale from *t* = 0, same as in the other conditions. (e) Measurement of primary reactive oxygen species (ROS) in the foldability measurement condition (see Methods for more details). The no-light and ROS-generating conditions serve as the negative and positive controls, respectively. TIr3 is an Ir-based photosensitizer generating O_2_^•–^ and ^1^O_2_ (Methods). The data are normalized by the negative controls and presented as mean ± SD (*n* = 3). One-way ANOVA with post-hoc Tukey HSD test (*** for *p* < 0.001, n.s. for *p* > 0.05). (f) Foldability trend over time under long-incubation conditions in a specific protein state. In the *U*-state condition, the *U* state is maintained at 35 pN for 90 min in every unfolding cycle. In the *N*-state condition, the *N* state is maintained at 1 pN for the initial 6 hr before the cyclic unfolding. (g) Proportion of water-exposed atoms in the *N* and *U* states for three different atom groups (mean ± SD).

We hypothesized that the accumulation of oxidative modification over time would cause the protein foldability decay. To test this hypothesis, we injected buffers with different oxidative stress levels after a few unfolding cycles (Fig. 3a, right) and assessed their impact on the protein foldability (Fig. 3c; Supplementary Fig. 1 for dissolved O_2_ concentration and pH of the buffers). Indeed, a vacuum-degassed or N_2_ gas-bubbled buffer with a reduced O_2_ concentration delayed the foldability decay with *τ* = ∼456 or 357 min, respectively. In contrast, when the protein was exposed to an O_2_ gas-bubbled buffer, the foldability was dramatically reduced to ∼53% even at the first unfolding after the buffer injection. 1% hydrogen peroxide (H_2_O_2_), a relatively mild ROS, also largely impaired the foldability, reducing it to ∼26% at the first unfolding, with an accelerated decay rate of *τ* = ∼22 min. We also found that the light irradiation facilitated the foldability decay. After cyclic unfolding for the initial 6 hr without the blue-light exposure, we observed a delay in the rate of foldability decay, similar to the case of the vacuum-degassed buffer (Fig. 3d). These results indicate that the foldability decay was primarily attributed to the photooxidative structural damage under blue light.

### Unusual mode of protein oxidative damage

In cellular and tissue environments, various biological oxidants like ROS are generated during metabolic reactions, such as the electron transport chain reaction and arginine metabolism^14,23–25^. However, no such reactions occur in our *in vitro* experiments. Furthermore, there is no exogenous photosensitizers, transition metal ions like Fe^2+^ essential for Fenton’s reaction, or high-energy ionizing radiation. As expected, the levels of the four primary ROS (H_2_O_2_, ^•^OH, O_2_^•–^, and ^1^O_2_) in our photoirradiation condition were comparable to those observed in the no-light condition (Fig. 3e). There may be a very low level of ROS, such as heat-generated H_2_O_2_ (ref. ^37^), although our experiments were conducted at room temperature (∼22°C), which is ∼20°C lower than in the previous report.

Under this condition of a low or negligible level of ROS, we observed an unexpected and remarkable finding: a rapid foldability decay even in the fully folded state (the *N* state). Despite maintaining the *N* state, which conceals numerous oxidizable atoms, at 1 pN for the initial 6 hr before cyclic unfolding, the foldability level still exhibited a significant decrease, similar to the control result with *τ* = ∼207 min (Fig. 3f). The foldability decrease in the *N*-state condition was even slightly faster than that of the *U*-state condition (*τ* = ∼293 min), where the fully unstructured polypeptide state (the *U* state) was maintained at 35 pN for a long time of 90 min at each unfolding cycle (∼95% of total experimental time at the *U* state). This result runs counter to the generally accepted notion, supported by previous experiments, that the compact, native state concealing a large portion of oxidizable atoms are less susceptible to oxidation compared to the unstructured, exposed states^21,22,38–41^. Indeed, the proportion of water-exposed, oxidizable atoms in the *N* state was estimated as 2–3 times lower than in the *U* state, regardless of analyzed atom groups (Fig. 3g and Extended Data Fig. 3). This unusual oxidative damage under the low or negligible ROS background appears puzzling, suggesting that there may be a hidden, unknown pathway of protein oxidation.

### Substantial oxidation near trapped O_2_ molecules within protein cavities

We conceived an alternative oxidation pathway that could account for the outcomes observed in the single-molecule tweezer experiments – dissolved O_2_ molecules become trapped within the intricate protein folds of the native state, promoting oxidative damage through close interactions within the diffusion-suppressing, protein core region. According to this oxidation pathway, amino acid residues inside the protein are expected to undergo more pronounced oxidation than those on the protein surface. To pinpoint the positions of oxidized residues, we performed liquid chromatography-tandem mass spectrometry (LC-MS/MS) analysis for blue light-exposed MBP (Methods and Extended Data Fig. 4a). From the volcano plot obtained from the LC-MS/MS analysis (Fig. 4a and Extended Data Fig. 4b,c), we identified 18 statistically significant oxidized residues under the blue-light condition (λ_peak_ = 446 nm; Extended Data Fig. 1 for the full spectrum), in comparison to the no-light condition (*p*-value < 0.05, fold change > 1.5; see Methods for more details). Indeed, the identified residues are predominantly located in the protein core region (Fig. 4b), accounting for 74.3% of their side chain atoms. This high proportion matches the prediction from the alternative oxidation scenario.

**Fig. 4.**
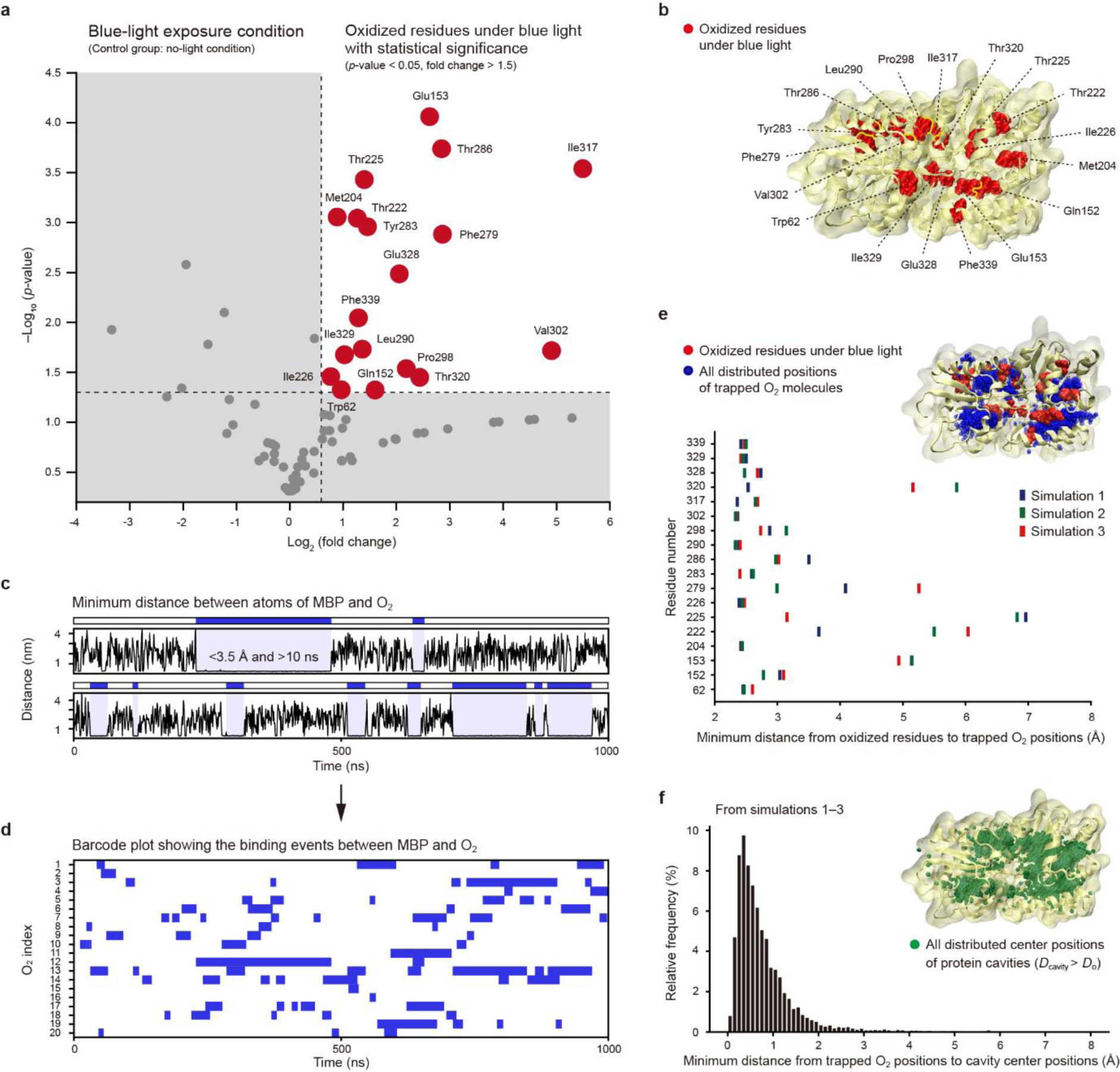
| Substantial oxidation near trapped O_2_ molecules within protein cavities. (a) Volcano plot showing significantly oxidized residues of maltose-binding protein (MBP) under blue light exposure (*p*-value < 0.05, fold change > 1.5; control: no-light condition). The amino acid residues are denoted as 3-letter code. The residue numbers indicate those following the signal peptide sequence. (b) Structural distribution of the oxidized residues. (c) Representative time traces of minimum distance between atoms of MBP and O_2_ from molecular dynamics (MD) simulations. The blue-colored regions indicate the bindings between the protein and O_2_, sustained within 3.5 Å for longer than 10 ns. (d) Barcode plot showing the binding events between MBP and all O_2_ molecules. (e) Minimum distances from oxidized residues to trapped O_2_ positions. The distances were calculated based on the positions of oxidized residues’ side chain atoms and trapped O_2_ molecules. The upper inset shows all distributed positions of trapped O_2_ molecules in simulation 1. (f) Minimum distances from trapped O_2_ positions to cavity center positions (only for protein cavities with *D*_cavity_ > *D*_o_). *D*_cavity_ and *D*_o_ represent the diameter of cavities and the kinetic diameter of O_2_ (3.46 Å), respectively. The upper inset shows all distributed center positions of the protein cavities with *D*_cavity_ > *D*_o_ in simulation 1.

We then turned to molecular dynamics (MD) simulations to investigate the potential trapping of O_2_ within the protein (Methods). From three 1-μs MD trajectories, we generated barcode-like plots that show the sustained and close bindings between MBP and O_2_ molecules (Fig. 4c,d). The blue-colored regions in the plots represent the binding time regions, where the minimum distance between their atoms remained below 3.5 Å for longer than 10 ns (see Methods for more details). We found that O_2_ positions during the binding times were predominantly distributed inside the protein with thermodynamic feasibility (Fig. 4e, upper inset; Extended Data Fig. 5a,b, Supplementary Methods, and Supplementary Video 1). Notably, the oxidized residues were in close proximity to the trapped O_2_ molecules, mostly within a range of 2.5–4.0 Å in their minimum distances (Fig. 4e). Using a tool called Voronoia that can identify protein cavities^42,43^ (Methods), we further examined whether the protein areas confining O_2_ molecules exhibited cavity-like structures. Indeed, the trapped O_2_ molecules were in close proximity to the center positions of protein cavities, mostly within a range of 0–2 Å in their minimum distances (Fig. 4f, Extended Data Fig. 5c,d, and Supplementary Fig. 2). Thus, the results of LC-MS/MS analysis, MD simulations, and protein cavity analysis collectively support the alternative oxidation pathway involving the trapping of O_2_ within the protein cavity spaces.

### Hydrophobic interactions as main driving force for O_2_ trapping

The primary regions where O_2_ molecules are trapped (Supplementary Figs. 3–5 and Supplementary Tables 1,2) are deeply located from the protein surface (Extended Data Fig. 6a,b). The predominant O_2_-trapping areas contain a large portion of hydrophobic residues, even more than the overall protein core region (Extended Data Fig. 6c). A hydropathy analysis shows that the O_2_-trapping areas exhibit larger hydrophobicity than nearly all successive sequence blocks of MBP (Extended Data Fig. 6d–f). This indicates that the distant residues in the sequence space would form the hydrophobic pockets for trapping O_2_. Additionally, the O_2_-trapping times were comparable to or even longer than the maltose-binding times, despite the function of MBP being the binding and transport of maltose into cells (Extended Data Fig. 7, Supplementary Figs. 6–9, and Supplementary Video 2). The results of these analyses suggest that the trapping events of nonpolar O_2_ molecules are mainly mediated by interactions with more hydrophobic regions of the cavity-comprising structural areas.

### Trp-mediated ^1^O_2_ generation under blue light

O_2_ is recognized to be kinetically stable^44^ and thus the direct oxidation by O_2_ would be unlikely to occur despite the local confinement and close contacts. It is noteworthy that under ultraviolet (UV) light, aromatic amino acids like Trp can function as endogenous photosensitizing species capable of generating a potent ROS, predominately ^1^O_2_, through the conventional energy transfer mechanism^26,27,45,46^. However, the range of direct UV absorbance of MBP, primarily through the aromatic residues, is entirely separated from our blue-light spectra in terms of wavelength and energy (λ_peak_ = ∼280 nm & *E*_peak_ = ∼4.4 eV *vs*. λ_peak_ = ∼450 nm & *E*_peak_ = ∼2.8 eV) (Extended Data Fig. 1). Intriguingly, a recent study has shown that when captured within ∼3 Å in a photosensitizing polymer network, O_2_ can be converted to ^1^O_2_ through a distinct mechanism involving spin-flip-based electron transfers^47^. Hence, using density functional theory (DFT) calculations, we examined whether the blue-light excitation can induce the generation of ^1^O_2_ through the spin-flip electron transfer mechanism and identified which amino acid residue is involved in this process. We then validated the ^1^O_2_ generation through spectrophotometric assays and electron paramagnetic resonance (EPR) spectroscopy.

We conducted mixed-reference spin-flip time-dependent DFT calculations^47,48^ (Methods), for each residue-O_2_ pair in four O_2_-trapping subareas exhibiting relatively tight O_2_ trapping for longer than 100 ns (Fig. 5a and Supplementary Fig. 4). In subarea 1, only the Trp62-O_2_ pair showed a noticeable absorption intensity at 445 nm (Fig. 5b), closely matching the peak wavelength of 447 nm used in our single-molecule tweezer experiments. The ground electronic state of the Trp62-O_2_ is the T_0_ state (^1^Trp62-^3^O_2_), with each *π*\* orbitals of O_2_ singly occupied. The two lowest triplet excited states (T_CT_; Trp62^•+^-O ^•–^) are nearly degenerate, corresponding to the charge transfer (CT) excitation from the Trp62 to the respective two *π*\* orbitals of O_2_ (Fig. 5c). The energy level of ∼2.7 eV for the T_0_→T_CT_ transition suggests that the Trp62-O_2_ can be excited in the blue-light spectrum. Additionally, the energies of the singlet excited states S_CT_ (∼2.7 eV) are nearly degenerate and very close to those of the T_CT_ states, indicating that an efficient intersystem crossing (ISC) can occur. The S_CT_ states can then undergo internal conversion to the S_3_ state (^1^Trp62-^1^O_2_(^1^Σ_g_^+^)) followed by the S_1_/S_2_ states (^1^Trp62-^1^O_2_(^1^Δ_g_)), ultimately resulting in ^1^O_2_ generation (Fig. 5d). The Trp129-O_2_ pair in subarea 3 also exhibited a similar absorption behavior to the Trp62-O_2_ pair in subarea 1, although the absorption intensity was relatively lower at 480 nm (Extended Data Fig. 8). These results suggest that the charge transfer excitation from Trp to O_2_ could be dominant in the blue-light spectrum (400–500 nm), compared to other types of residues.

**Fig. 5.**
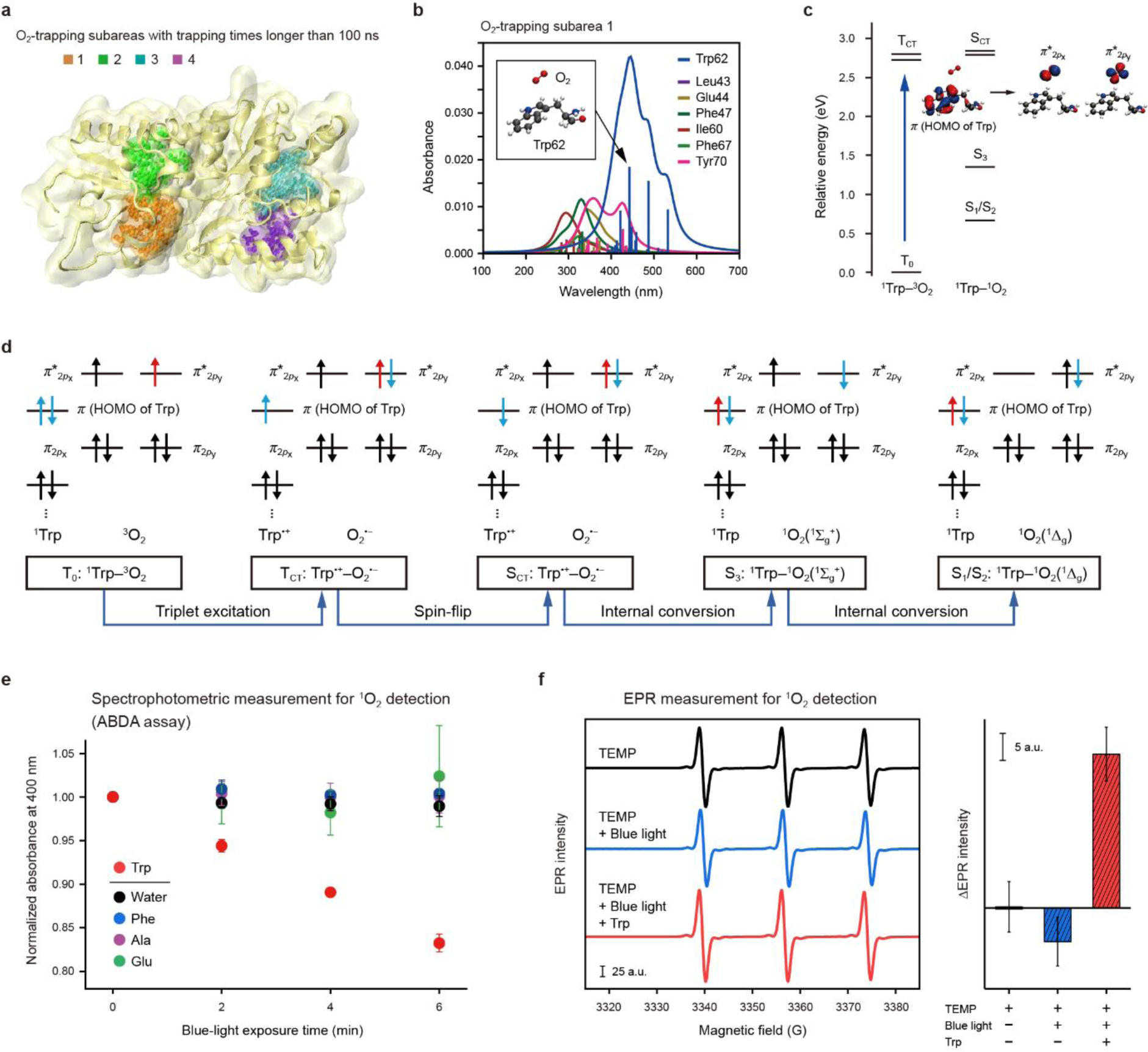
| Trp-mediated ^1^O_2_ generation under blue light. (a) O_2_-trapping subareas exhibiting relatively tight O_2_ binding for longer than 100 ns. In each subarea, the amino acid residues highlighted in color are those whose atoms are within 3.5 Å from O_2_ atoms during the binding periods. (b) Absorption spectra of residue-O_2_ pairs in O_2_-trapping subarea 1 from density functional theory (DFT) calculations. Smooth curves were obtained by Lorentzian broadening from spectral peaks with a width of 40 nm. The residues are denoted as 3-letter code. The residue numbers indicate those following the signal peptide sequence. The inset shows the Trp62-O_2_ structure with the maximum oscillator strength. (c) Relative energy diagram for singlet and triplet states of Trp62-O_2_ pair. The molecular orbitals are shown for the transition from T_0_ to T_CT_ state. (d) Molecular orbital diagrams and overall pathway for ^1^O_2_ generation. (e) Spectrophotometric measurement for ^1^O_2_ detection using ABDA assay. The reaction of ABDA with ^1^O_2_ decreases the absorbance. Water indicates deionized (DI) water. (f) Electron paramagnetic resonance (EPR) measurement for ^1^O_2_ detection. The EPR spectrum and intensity change for each condition are shown on the left and right, respectively. The reaction of TEMP with ^1^O_2_ increases the EPR intensity.

Using spectrophotometric assays and EPR measurements (see Methods for more details and the full names of molecules), we confirmed the Trp-mediated ^1^O_2_ generation when exposed to blue light. For a Trp solution containing ABDA as a ^1^O_2_ probe, we observed a gradual decrease in absorbance at 400 nm during the blue-light exposure, indicating the ^1^O_2_-induced ABDA oxidation (Fig. 5e and Extended Data Fig. 9). In contrast, other tested amino acids (aromatic: Phe, charged: Glu, hydrophobic: Ala) did not exhibit noticeable changes in absorbance (Fig. 5e and Extended Data Fig. 9). Additionally, in the EPR measurements using TEMP (a commonly used ^1^O_2_ spin trapper), we observed an increase in the EPR intensity in the Trp/light condition, due to the formation of the TEMPO radical (the oxidized form of TEMP), while a decrease occurred in the condition of only blue-light exposure (Fig. 5f). The decrease in EPR intensity may result from the light-induced degradation of the oxidized product. However, the opposite result in the Trp/light condition indicates the formation of the oxidized product in sufficient amounts. The spectrophotometric/EPR measurements and DFT calculations collectively support the blue light-induced ^1^O_2_ generation near Trp residues within the protein.

### Close proximity between Trp and oxidized residues

Considering the Trp-mediated ^1^O_2_ generation, the oxidized residues detected in the LC-MS/MS analysis are expected to be in close proximity to Trp residues. Despite the apparent proximity of Trp residues to the oxidized residues (Fig. 6a), we further conducted Monte Carlo (MC) analysis to quantitatively assess this proximity (see Methods for more details). After randomly sampling the residues from each type, we calculated the average distances between the oxidized residues and sampled residues over all possible combinations in each random sampling. This process was iterated to construct a probability density distribution for each residue type. Remarkably, Trp emerged as the closest residue among all types of residues (Fig. 6b and Extended Data Fig. 10).

**Fig. 6.**
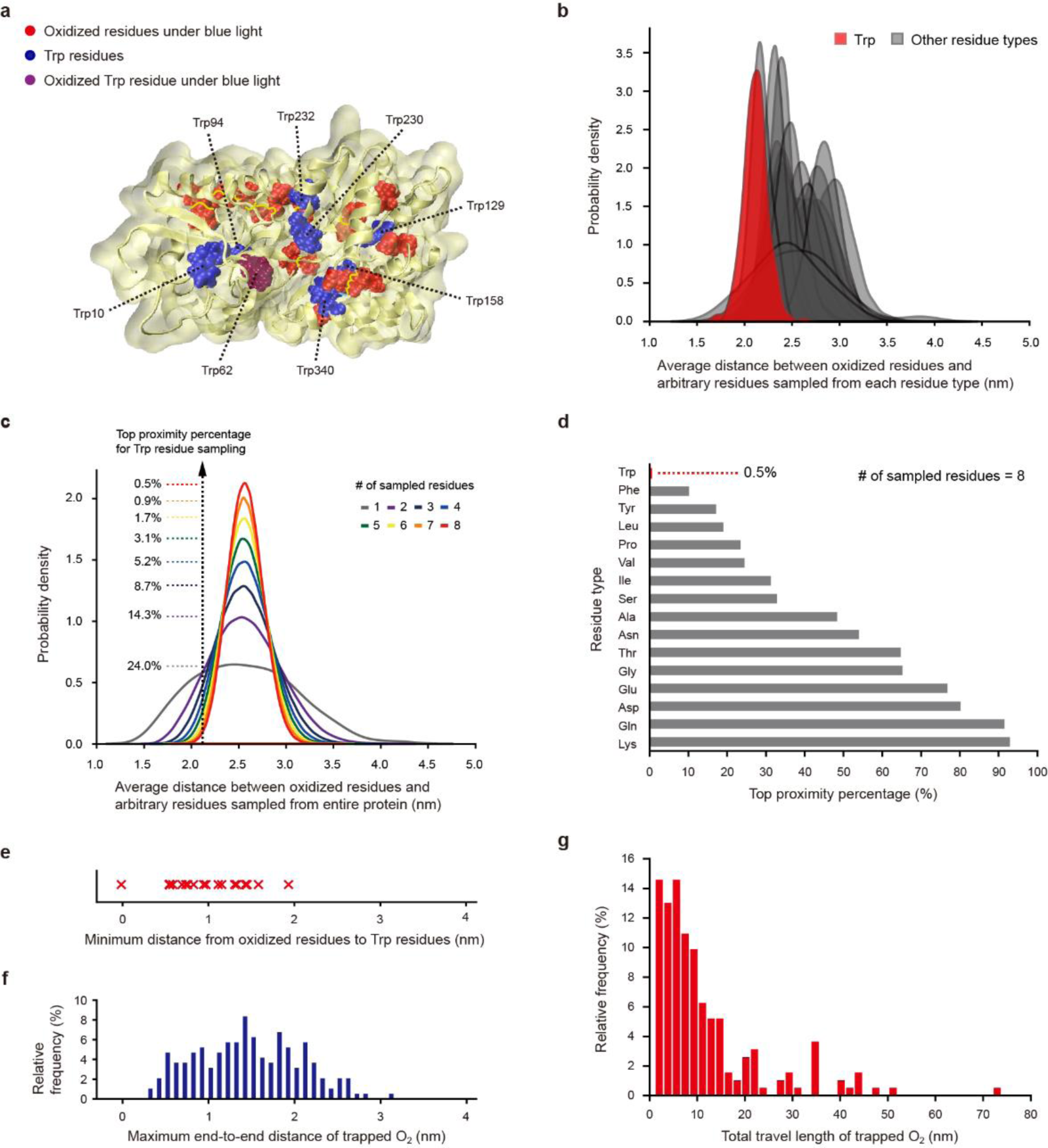
| Close proximity between Trp and oxidized residues. (a) Structural distribution of all tryptophan (Trp) residues and oxidized residues. The residue numbers indicate those following the signal peptide sequence. (b–d) Monte Carlo (MC) analysis for the proximity between Trp and oxidized residues (see Methods for more details). The data for other residue types in panel b are distinguishably presented in Extended Data Fig. 10. The top proximity percentages in panel c are shown for the cases with different numbers of sampled residues up to the total number of Trp residues (*n* = 8). The panel d shows the histogram of the top proximity percentages for all other residue types with a larger number of residues than the total number of the benchmark Trp residues, in ascending order. The residue proportions are shown in Extended Data Fig. 6c. The residues are denoted as 3-letter code. (e) Minimum distances from oxidized residues to Trp residues. (f,g) Maximum end-to-end distances (f) and total travel lengths (g) of trapped O_2_ molecules within the protein obtained from molecular dynamics (MD) simulations.

To see how likely this close proximity occurs, we compared the average distances analyzed for Trp with those between the oxidized residues and arbitrary residues sampled from the entire protein (Fig. 6c). As the number of sampled Trp residues increased, the proximity approached the top 0.5% in the probability density distribution constructed for any residue sampling (Fig. 6c). Hydrophobic residues tend to show relatively high proximity (*i.e.*, low percentage values) due to their prevalence in the protein core region, where the oxidized residues are mainly located (Fig. 6d). However, their proximity is limited to the top 10–50%, with Trp being the notable exception (Fig. 6d). This analysis implies that the close proximity between the oxidized residues and Trp residues is less likely to occur by chance. It indicates a probable correlation in their positions, reconciling the Trp-mediated ^1^O_2_ generation and subsequent oxidation of nearby residues.

The individual oxidized residues from Trp residues are all less than 2 nm in their minimum distances (Fig. 6e). This distance range falls within the maximum end-to-end distances and total travel lengths of trapped O_2_, reaching up to ∼3 nm and tens of nm, respectively (Fig. 6f,g and Supplementary Video 1). These estimated distance ranges for constrained O_2_ diffusion would also be applicable to ^1^O_2_ for two main reasons, if no reaction occurs en route: (1) hydrophobic interactions are likely the primary driving force behind the O_2_ trapping, as discussed earlier, and (2) the overall polarity would remain constant after the conversion to ^1^O_2_ due to the symmetrical distribution of both *π*\* orbitals over the two oxygen atoms (Fig. 5c). Moreover, the free diffusion of ^1^O_2_ was reported to be very long (∼1 cm in air and ∼125 nm in water^49,50^). However, in contrast to the free diffusion, the rugged structural shapes of local areas would restrict the paths of the constrained diffusion and modulate the reaction rates for individual residues^41,51^, possibly resulting in various types and dispersed locations of the oxidized residues observed in the LC-MS/MS analysis.

## Discussion

Synthesizing all our results and findings, we propose an O_2_-confinement pathway for protein oxidative damage, comprising four main steps (Fig. 1): (1) the confinement of O_2_ within protein cavities, (2) the photoexcitation of Trp-O_2_ pair system under blue light (400–500 nm), (3) the generation of ^1^O_2_ through the spin-flip electron transfer, and (4) the oxidation of nearby residues by ^1^O_2_ through constrained diffusion within the protein.

According to this oxidation mechanism, protein folds capable of capturing O_2_ could be inherently more vulnerable to oxidative damage. Due to the thermal fluctuation of structural conformations, some proteins may experience more frequent and more broadly distributed trapping of O_2_ molecules than expected for their static structures. Given that even a single mutation can critically impair the protein stability/fold^52^, a single oxidative modification at one residue could severely impact the protein integrity. Thus, the oxidative damage to proteins with numerous cavity spaces, such as MBP we studied here, could be rapidly accelerated, since they can capture O_2_ molecules at multiple sites, thereby enhancing the potential for oxidative attacks. Taken together, the O_2_-confinement oxidation we describe here would be a crucial factor to consider in maintaining protein stability and function, especially under *in vitro* conditions with a high pO_2_ level of ∼21% and easy exposure to visible light. Our study suggests that the inherent susceptibility to oxidative damage is partially encoded in the internal architecture of protein folds, highlighting key elements to consider when designing more robust proteins resistant to oxidation. It is also advisable to avoid using blue light in bioimaging methods whenever possible to prevent the potential facilitation of protein oxidative damage.

It has been reported that the blue-light irradiation can directly or indirectly lead to the tissue aging and pathological states, particularly in visible light-penetrating tissues, such as skin and eyes^53–58^. This association has been suggested to primarily arise from an elevated oxidative stress caused by the blue light-induced ROS generation through endogenous photosensitizers like flavins^53,59–63^. The pO_2_ levels in the skin/eye tissues are not particularly substantial compared to other tissues, but rather in hypoxic to moderate conditions^28,64,65^. Their antioxidant defense systems^66–68^, which target freely-diffusing ROS, will further mitigate the oxidative stress. Remarkably, the O_2_-confinement oxidation described here can attack proteins by evading the antioxidant networks through the initial O_2_ trapping. Thus, through this alternative photooxidation pathway, even a low level of dissolved O_2_ can more easily compromise the integrity of proteins. Additionally, enzymatic repair of this type of oxidative damage would be limited due to the inaccessibility to the attacked loci buried within the protein. Therefore, the O_2_-confinement oxidation may constitute a fundamental and key oxidation layer contributing to the overall protein damage, aging, and disorders in the skin/eye tissues, especially given that modern lifestyles increasingly rely on LED lighting and electronic displays emitting a high amount of blue light (ref. ^57^).

## Methods

### Protein expression and purification

We studied maltose-binding protein (MBP) without the signal peptide sequence, as in previous single-molecule forced unfolding studies^33–36^ (Extended Data Fig. 2a for the amino acid sequence). The SpyTag, a short peptide tag, was introduced at the N- and C-terminal ends for the DNA handle attachment. The corresponding gene block was cloned into pET24a vector. The pET24a vectors were inserted into competent cells (*E. coli* BL21(DE3)) using heat shock transformation. The cells were cultured in 1 L of LB medium with kanamycin (25 µg/mL) at 37°C. Around OD_600_ = 1.0, the protein was overexpressed by 0.8 mM IPTG for 4 hr at 30°C. The cells were pelleted by centrifugation (5700 rpm, 10 min, 4°C), resuspended in 50 mM Tris-HCl, pH 7.4, 200 mM NaCl, 1 mM TCEP, 10% glycerol, 1 mM PMSF, and then lysed using Emulsiflex C3 at ∼17000 psi. After another round of centrifugation (18000 rpm, 30 min, 4°C), 1 ml Ni-IDA resin was added to the supernatant and incubated for 1 hr. The resin was washed with 50 mM Tris-HCl, pH 7.4, 200 mM NaCl, 1 mM TCEP, 10% glycerol, 20 mM imidazole. The protein was eluted from the resin with 50 mM Tris-HCl, pH 7.4, 200 mM NaCl, 1 mM TCEP, 10% glycerol, 300 mM imidazole. The eluted protein was stored at –80°C in aliquots.

### DNA handle conjugation to protein

1022-bp DNA handles were attached to MBP for the single-molecule forced unfolding experiments^31,69^. The DNA constructs, modified at one end with an amine group and at the other end with either azide or 2×biotin, were generated by PCR using λ DNA template (NEB, N3011S)^31^. ∼8 ml PCR product (azide-DNA:2xbiotin-DNA = 1:1) was purified using HiSpeed Plasmid Maxi kit (Qiagen) in 1 ml NaHCO_3_ (pH 8.3). The amine group of the purified DNA constructs was further modified to maleimide by incubating the sample with 1 mM SM(PEG)_2_ (Thermo Scientific Pierce) for 20 min at ∼23°C. The sample was subsequently purified using Econo-Pac 10DG Desalting Column (Bio-Rad) in 1.5 ml of 0.1 M sodium phosphate (pH 7.3) with 150 mM NaCl. ∼0.5 µM of the DNA sample were incubated with ∼20 µM of a SpyCatcher construct for 2 hr at ∼23°C (refs. ^31,69^), for the molecular conjugation through the functional groups of maleimide and cysteine thiol. Unconjugated proteins were removed using anion exchange chromatography (HiTrap Q HP column, Cytiva) with a gradient mode of 0–1 M NaCl in 20 mM Tris-HCl (pH 7.5). The DNA peak fractions, ∼30% of which corresponds to SpyCatcher-conjugated DNA, was concentrated to ∼100 nM for the conjugated construct. 1 µl of ∼20 µM MBP was incubated with 10 µl of the DNA sample for 2 hr at ∼23°C, for the DNA handle attachment to MBP at the N- and C-terminal ends through the SpyTag-SpyCather binding. A covalent, isopeptide bond is spontaneously formed in the bound complex^70^ and its unfolding is not observed in our force range up to 50 pN (refs. ^31,69^). The final sample was diluted to make the hybrid molecular construct with azide and 2xbiotin at both termini to be ∼200 pM and stored at –80°C in aliquots.

### Preparation of single-molecule sample chambers

The surfaces of coverslips (VWR, No. 1.5, 24×50 and 24×40 mm) were cleaned using KOH and Piranha solution^31,69^. The bottom coverslip (24×50 mm) was passivated with polyethylene glycol (PEG) polymers of methyl-PEG and biotin-PEG at a molar ratio of 100:1 (refs. ^31,69^). These two surface-treated coverslips were combined to construct a single-molecule sample chamber^31,69^, with a channel volume (CV) of ∼10 µl (≡ 1 CV). 1 µl of polystyrene beads coated with streptavidin was washed with 0.1 M sodium phosphate, pH 7.4, 150 mM NaCl, 0.1% Tween 20 (ref. ^31^). 1 CV of the bead sample was injected into the sample chamber and incubated for 2–5 min at ∼22°C for the surface binding. The polystyrene beads bound to the surface were used to correct the thermal drift of the microscope stage. 100 mg/ml of bovine serum albumin (BSA) was injected into the chamber and incubated for 5 min at ∼22°C for additional surface passivation. The chamber was washed with 50 mM Tris-HCl, pH 7.4, 150 mM NaCl (Buffer A). 10 µl of ∼200 pM MBP-DNA construct sample was mixed with 1 µl of 0.2 µM traptavidin (TTV) for 15 min at ∼22°C. 1 CV of the MBP-DNA-TTV construct sample (100–200 pM) was injected into the chamber and incubated for 5 min at ∼22°C for the surface binding. To block unoccupied biotin-binding sites of TTV, 1 CV of a biotin-labeled oligonucleotide (30 nucleotides; 10 µM in Buffer A) was injected into the chamber and incubated for 5 min at ∼22°C. The chamber was washed again with Buffer A. 1 µl of DBCO-coated magnetic beads was washed and resuspended with Buffer A (ref. ^31^). 1 CV of the bead sample was injected into the chamber and incubated for 1 hr at ∼25°C. The beads are covalently attached to the surface-tethered molecular constructs through dibenzocyclooctyne (DBCO)-azide conjugation. The sample chamber was finally washed with 50 mM HEPES, pH 7.6, 100 mM KCl, 5 mM MgCl_2_ (Buffer B), which was used in previous single-molecule forced unfolding studies on MBP^33,34,36^.

### Single-molecule forced unfolding experiments

The single-molecule forced unfolding experiments were conducted at ∼22°C using a custom-built magnetic tweezer apparatus^31,71^. The assembled sample chamber was placed on an inverted microscope (Olympus, IX73) equipped with a motorized sample stage (ASI, MS-2000 XY Automated Stage). The imaging spots were illuminated using a blue light-emitting diode (LED; Thorlabs, M455L4, λ_peak_ = 447 nm; Extended Data Fig. 1 for the full spectrum). The force applied to a tethered magnetic bead was pre-calibrated using an inverted pendulum model of the bead-molecular construct^31,72^. The extension (end-to-end distance) of the protein-DNA construct was determined by tracking changes in bead diffraction patterns captured by a charge-coupled device (CCD) camera (JAI, CM-040 GE)^31,72^. For the control experiments, the force-scanning range for each unfolding cycle was 1–50 pN with waiting times between the cycles as 2.5 min at 1 pN and 1 s at 50 pN. Especially for the experiments involving a prolonged incubation in the *U* state, the maximum force was set to 35 pN for 90 min in every unfolding cycle. This force level is sufficiently high to induce the unfolding and is relatively more resistant to bead detachment during the waiting time. 93% were fully unfolded during the stretching phase before reaching 35 pN, whereas 7% were fully unfolded at 35 pN within 0.66±0.16 s (mean ± SD). The total incubation time for the *U* state was estimated to be ∼95% of the total experiment time. In the *N*-state condition, the *N* state was maintained at 1 pN for the initial 6 hr before the cyclic unfolding. The foldability curves in Fig. 3 were smoothed using a moving-average window of three data points to reduce fluctuation noise and obtain more accurate decay time constants. The curves for the control condition and other conditions of the same large number of data points per unit time were binned into groups of five data points for clearer visualization and comparison.

### Analysis of force-extension curve data

The force-extension curves were median-filtered for the extension (half-window size = 5) and smoothed for the force (half-window size = 10). The unfolding forces and step sizes extracted from the force-extension curves were analyzed using the worm-like chain (WLC) model^73^, *FL*_p_/*k*_B_*T* = *l*/*L* + (1–*l*/*L*)^−2^/4 – 1/4, where *F* is the applied force, *k*_B_ is the Boltzmann constant, *T* is the absolute temperature, *l* is the protein extension, *L*_p_ is the persistence length, and *L* is the contour length. The contour length was estimated as the number of amino acid residues multiplied by the average residue-residue distance of 0.38 nm (ref. ^71^), followed by correction with protein structure factors. The persistence length was estimated as 0.29±0.01 (SE) nm for the external α-helices at the C-terminus and 0.51±0.01 (SE) nm for the fully unstructured polypeptide from the WLC model, which aligns with a range of protein persistence lengths^74,75^. The slight variance in persistence length for each case is likely due to the difference in residue composition within the unstructured portions^74,76^. The most probable unfolding forces for the external α-helices and the core structure were measured as 13.9±0.2 pN and 25.3±0.7 pN (peak value ± SE), respectively. The foldability at each unfolding cycle was estimated by the proportion of normal unfolding with the distinctive pattern, *i.e.*, the unfolding of the external α-helices followed by the core structure, obtained from multiple molecules.

### Gas sparging and vacuum degassing

A vial containing 5 ml Buffer B was sealed with a rubber septum and parafilm to block air flow. For the gas sparging, a gasbag filled with either nitrogen or oxygen gas was connected to a 2 ml syringe with a needle. The needle linked to the gasbag was inserted into the buffer solution through the septum/parafilm. Another needle was inserted into the air above the buffer through the septum/parafilm, allowing the gas from the gasbag to replace the dissolved gas by the pressure difference. The gas sparging process was maintained at ∼22°C for 30 min. For the vacuum degassing, a vial containing 5 ml Buffer B was connected to a vacuum pump for 1 hr. To facilitate the degassing, the vial was rapidly frozen using liquid nitrogen and then thawed during the vacuum degassing. The freezing and thawing was repeated three times. After the gas sparging or vacuum degassing, the concentration of dissolved molecular oxygen (O_2_) and buffer pH were immediately measured (Supplementary Fig. 1).

### Measurement of reactive oxygen species (ROS)

The levels of four primary ROS (hydrogen peroxide (H_2_O_2_), hydroxyl radical (^•^OH), superoxide (O_2_^•–^), singlet oxygen (^1^O_2_)) were measured after 10 min incubation at ∼22°C in three conditions: the foldability measurement condition (Buffer B with 0.12 mg/ml magnetic beads under blue light), negative control (no-light condition), and positive control (ROS-generating condition). Absorbance or fluorescence was obtained as a signal from 200 µl samples using a microplate reader. The values for the three conditions were normalized by that of the negative control. Peroxidase (POD)-catalyzed oxidation of *N*,*N*-diethyl-*p*-phenylenediamine (DPD) by H_2_O_2_ was measured for the assessment of H_2_O_2_ level (DPD assay)^77^. 1 mM H_2_O_2_ was used for the positive control. After 10 min incubation, each 200 μl sample was mixed with 80 μl of 0.1 M sodium phosphate (pH 6.0), 224 μl deionized (DI) water, 10 μl of 1% w/v DPD (Merck, 07672), and 10 μl of 0.1% w/v POD (Thermo Fisher, 31490). The H_2_O_2_-induced generation of the radical cation of DPD (DPD^•+^) was manifested by an increase at OD_510_ (ref. ^77^). Oxidation of hydroxyphenyl fluorescein (HPF) by ^•^OH was measured for the assessment of ^•^OH level (HPF assay)^78^. 5 μM HPF (Cayman Chemical, 10159) was dissolved in each buffer solution. A Fenton’s reagent (100 μM FeSO_4_, 1 mM EDTA, 0.3% H_2_O_2_) was used for the positive control. The fluorescence from oxidized HPF by ^•^OH was measured with λ_ex_ = 485 nm and λ_em_ = 525 nm (refs. ^47,79^). Oxidation of dihydrorhodamine 123 (DHR123) by O_2_^•–^ was measured for the assessment of O_2_^•–^ level (DHR123 assay)^47,80^. 5 μM DHR123 (Thermo Fisher, D23806) was dissolved in each buffer solution. 0.5 μM of an iridium-complex photosensitizer (TIr3) synthesized as previously described^80^ was used for the positive control. TIr3 was reported to generate O_2_^•–^ under a blue-light irradiation^80^. The fluorescence from the oxidized DHR123 (rhodamine 123) by O_2_^•–^ was measured with λ_ex_ = 485 nm and λ_em_ = 525 nm (ref. ^81^). Oxidation of 9,10-anthracenediylbis(methylene)dimalonic acid (ABDA) by ^1^O_2_ was measured for the assessment of ^1^O_2_ level (ABDA assay)^82^. 100 μM ABDA (Merck, 75068) was dissolved in each buffer solution. 0.5 μM TIr3 was used for the positive control as TIr3 was also reported to generate ^1^O_2_ under a blue-light irradiation^80^. ^1^O_2_-induced ABDA oxidation was manifested by a decease at OD_400_ (refs. ^80,82^). The absorbance for the positive control was corrected by that without ABDA since TIr3 itself has absorbance around 400 nm (ref. ^80^). The ABDA assay was also employed to detect ^1^O_2_ in solutions with a high concentration of free amino acids (refer to the caption of Extended Data Fig. 9 for more experimental details).

### Sample preparation for LC-MS/MS

The procedure of sample preparation for liquid chromatography-tandem mass spectrometry (LC-MS/MS) was modified from previous studies^83,84^. 10 µg MBP was added to either the same volume of DI water upon O_2_- or Ar-bubbling. For control groups not exposed to light, the protein samples in the O_2_- or Ar-bubbled condition was kept in a dark chamber for 30 min. Each group corresponds to a high or low concentration of dissolved O_2_ (Supplementary Fig. 1) and show little difference in oxidation based on a volcano plot (Extended Data Fig. 4). For the experimental group subjected to light exposure, the protein sample in the O_2_-bubbled condition was irradiated with a blue LED (λ_peak_ = 446 nm, Extended Data Fig. 1 for the full spectrum) for 30 min. Every sample was further purified using SDS-free polyacrylamide gel electrophoresis (native PAGE) for LC-MS/MS, which does not induce protein denaturation. To prevent undesirable protein oxidation caused by ammonium persulfate during gel running, 0.01% (v/v) thioglycolic acid, an antioxidant, was added to sample loading buffer^85,86^. N_2_-bubbled DI water was used in making a 7.5% SDS-free bis-acrylamide gel and gel-running buffer. The electrophoresis was carried out for 1.5 hr at 160 V, maintaining a temperature of 4°C. Following washing with LC-grade water, the gel was stained using Imperial^TM^ Protein Stain (Thermo Sicentific, 24617). After destaining with LC-grade water, the gel section containing the major band was cut into ∼1 mm^3^ cubes using a razor blade and transferred to a non-treated 96-well plate. The gel cubes were further destained with MeOH, 50 mM ammonium bicarbonate (ABC), and 40% acetonitrile (ACN) (aq), and then dehydrated with ACN. Extra washing and dehydration steps were carried out with LC-grade water, 40% ACN (aq), and ACN. 1 µg/µl trypsin in MS-grade acetic acid was diluted to 0.01 µg/µl using a 9:1 (v/v) mixture of 50 mM ABC and ACN (Buffer X). 25 µl of the trypsin solution and 75 µl of Buffer X were added to the dehydrated gel cubes. After a 5-min wait, the 96-well plate was sealed with parafilm and incubated for ∼18 hr at 37°C. The resulting tryptic peptides of MBP were eluted from the gel cubes by 0.1% formic acid/50% ACN (aq), and then transferred to Eppendorf Protein LoBind tubes. Additional elution was performed using 0.1% formic acid/99.9% ACN. The eluted samples were dried using SpeedVac at 60°C, resuspended in 0.1% trifluoroacetic acid (TFA) (aq), and subjected to clean-up using Pierce^TM^ C18 Tips (Thermo Scientific, 87782). The tips were wet and then equilibrated by aspirating and discarding 50% ACN (aq) followed by 0.1% TFA (aq). The peptide suspension was slowly and repetitively aspirated and dispensed (×10). After rinsing the tip with 0.1% TFA/5% ACN (aq), the peptides were eluted by slowly aspirating and dispensing 0.1% formic acid/80% ACN (aq). The eluted peptides were transferred to fresh LoBind tubes, dried using SpeedVac at 60°C, and resuspended in 0.1% formic acid (aq) for LC-MS/MS analysis. All experiments were conducted in triplicate.

### LC-MS/MS analysis

After the in-gel tryptic digestion, the resulting tryptic peptides were analyzed by LC-MS/MS. The tryptic digest was separated through online reversed-phase chromatography using a peptide trap column Acclaim PepMap 100 C18 (Thermo Fisher Scientific; 3 μm particle size, 75 μm diameter, 2 cm length) and an analytical column Acclaim PepMap RSLC C18 (Thermo Fisher Scientific; 3 μm particle size, 75 μm diameter, 15 cm length). The procedure was followed by electrospray ionization at a flow rate of 300 nl/min. The samples were eluted using a gradient of 3–50% solution B (80% ACN/0.1% FA (aq)) for 60 min and 50–80% solution B for 10 min, and then the columns were washed with 100% solution B for 10 min. The chromatography system was coupled in-line with an Orbitrap Fusion Lumos Tribrid mass spectrometer (Thermo Fisher Scientific). The mass spectrometer was operated in the data-dependent mode with a 120,000-resolution MS1 scan (375–1500 m/z), an AGC target of 5e5, and a maximum injection time of 50 ms. Peptides above a threshold of 5e3 and with a charge of 2–7 were selected for fragmentation with dynamic exclusion. The dynamic exclusion duration was set to be 15 s with 10-ppm mass tolerance.

### LC-MS/MS data processing

All MS/MS spectra were analyzed using SEQUEST (Thermo Fisher Scientific; version IseNode in Proteome Discoverer 2.4.1.15), matching against rcsb_pdb_1ANF.fasta for MBP with following parameters: 10 ppm for parent mass tolerance; 0.6 Da for fragment mass tolerance; fully digested by trypsin; fixed modification: carbamidomethylation (cysteine); variable modifications: Met-loss/Met-loss+acetylation (N-terminal methionine), deamidation (asparagine and glutamine), acetylation (lysine and N-terminus), phosphorylation (serine, threonine, and tyrosine), oxidation (arginine, asparagine, aspartic acid, glutamic acid, glutamine, histidine, isoleucine, leucine, lysine, methionine, phenylalanine, proline, serine, threonine, tryptophan, tyrosine, and valine; +15.99 Da resulting from oxygen addition or hydroxylation), based on the UNIMOD database. Cysteine was excluded from the variable modification of the oxidation due to its absence in MBP. The peptide and protein identifications were validated using Scaffold Q+ (Proteome Software; version 5.2.2). Identified peptides were accepted when >95% probability was obtained from Scaffold Local FDR algorithm. Identified proteins were accepted when >95% probability was obtained from Protein Prophet algorithm and at least two identified peptides were contained. The oxidation level for each amino acid residue was quantified as the oxidative modification count for the residue divided by the total detected peptide count containing the residue. All the LC-MS/MS experiments and analysis were performed in triplicate for each control and experimental group. The volcano plots were generated to compare the oxidation level between two groups. In the volcano plots, the fold change values were obtained based on the mean oxidation level for each residue from the triplicate data. Residues of no detection or zero oxidation in all triplicate data were omitted in the volcano plots because the fold change cannot be defined. Significantly oxidized residues compared to the control group were selected as those with *p*-value < 0.05 and fold change > 1.5 (ref. ^87,88^).

### Molecular dynamics (MD) simulations

All MD trajectories were generated using CHARMM36m force field (charmm36-mar2019)^89^, which were implemented in the GROMACS molecular dynamics software package (version 2021.4). The input files of simulation systems were generated using CHARMM-GUI^90,91^. The initial coordinates for the same MBP construct used in the foldability measurements were obtained from a crystal structure (PDB ID 1ANF)^92^. The systems were solvated with TIP3P water molecules^93^. K^+^ and Cl^−^ ions were added to neutralize the systems and maintained at ∼150 mM. The force field and structure files of O_2_ molecule were generated using CHARMM-GUI Ligand Reader & Modeler^94^. The force field and structure files of maltose was generated using Open Babel^95^ and SwissParam^96^. Using a Gromacs module of gmx insert-molecules, twenty O_2_ or one maltose molecule were inserted into the systems. Total numbers of atoms were 61600–61700 for all the simulation systems. The van der Waals (vdW) interactions were calculated with a cutoff distance of 12 Å and smoothly switched off at 10–12 Å by a force-switch function^97^. The long-range electrostatic interactions were calculated using the particle-mesh Ewald method with a cutoff distance of 12 Å and a mesh size of 1.2 Å (ref. ^98^). During the equilibration run, the NVT ensemble was applied with 1-fs time step for 125 ps. The temperature was maintained at 303.15 K using the Nosé–Hoover temperature coupling method with *τ*_t_ = 1 ps (refs. ^99,100^). The production run was performed with 2-fs time step in the NPT ensemble. The pressure was maintained at 1 bar using the isotropic Parrinello-Rahman method with *τ*_p_ = 5 ps and a compressibility of 4.5×10^−5^ bar^−1^ (refs. ^101,102^). The production times were 1 μs for the MBP/O_2_ and MBP-only systems and 15 μs for the MBP/maltose system. The MD simulation trajectory data were analyzed using Gromacs modules and custom-built Python codes. After the centering of MBP in the MD simulation systems, the translational and rotational movements of the protein were eliminated in the MD trajectories using ‘-fit rot+trans’ option of gmx trjconv module. Visual Molecular Dynamics (VMD) was used for the graphical visualization and movie generation^103^.

### Estimation of protein surface and core atoms

The solvent-accessible surface (SAS) of MBP was analyzed using Gromacs solvent-accessible surface area (SASA) module. The SAS was generated with a probe radius (*r*_probe_) of 1.4 Å in every 100 ns of a MD simulation. 1.4 Å is a typically used value for *r*_probe_, which approximates the radius of a water molecule. The distances between the atom centers and SAS (*d*_atom-SAS_) were calculated for all protein atoms. By subtracting the vdW radius (*r*_vdW_)^104^ and the probe radius from *d*_atom-SAS_, the distances between the vdW surface of atoms and the solvent-excluded surface (SES) (*d*_vdW-SES_) were calculated for all protein atoms. The relative frequencies of *d*_vdW-SES_ were calculated for all protein atoms, all side-chain atoms, and highly oxidizable side-chain atoms^105^. For each case, the protein surface or core atoms were classified based on *d*_vdW-SES_ values smaller or larger than 0.1 Å, respectively (Extended Data Fig. 3).

### Position and distance analyses of trapped O_2_ molecules and protein cavities

The minimum distance between atoms of MBP and O_2_ molecules was extracted from the MD trajectories in every 1 ns. We collected all time points, during which the distance remains less than a cutoff distance for longer than a cut-off time span. 3.5 Å was used as the cutoff distance to include only the close binding. The cutoff distance value of 3.5 Å has been used to determine the atoms involved in protein-ligand interactions^106–108^. 10 ns was used as the cutoff time span to exclude a large number of transient interactions and only consider more sustained binding events. In this analysis, the first and the last time frames for each duration were excluded to analyze the time regions of complete O_2_ trapping within the protein. The positions of O_2_ molecules as a center coordinate of O_2_ atoms were extracted from all the collected time points and superimposed onto the MBP structure. The maximum end-to-end distances and total travel lengths of individual trapped O_2_ molecules were also calculated for all O_2_-trapping time regions. For the protein cavity analysis, the atom coordinates of MBP were extracted as PDB files from the MD trajectories in every 1 ns. For every PDB file, the center positions and diameters of protein cavities were obtained using Voronoia with a grid distance of 0.2 Å (refs. ^42,43^). We only considered the cavities with a diameter larger than the kinetic diameter of O_2_ (3.46 Å). The center positions of the protein cavities were superimposed onto the MBP structure.

### Depth and hydrophobicity analyses for predominant O_2_-trapping (sub)areas

We first determined the predominant O_2_-trapping (sub)areas through the procedures described in Supplementary Methods. The depths of the O_2_-trapping (sub)areas from the protein surface (solvent-excluded surface) were calculated from three MD simulations in every 100 ns. The distances between the protein surface and atom centers (atom depths) were calculated using Michel Sanner’s Molecular Surface (MSMS) for each time point^109^. The atom depths were averaged for all amino acid residues comprising the O_2_-trapping (sub)areas (area depths). The area depths were further averaged over the analyzed time points. The Moon-Fleming hydrophobicity scale for amino acid residues^110^ was utilized in the hydrophobicity analyses for the predominant O_2_-trapping (sub)areas. The hydrophobicity scale is expressed as the free energy difference in stability (ΔΔ*G*) between the wild-type transmembrane (TM) protein OmpLA and each amino acid variant at the position A210 (*e.g.*, for cysteine, ΔΔ*G* = Δ*G*_WT_– Δ*G*_A210C_). In this scale, more hydrophobic amino acid has more negative ΔΔ*G*. The hydrophobicity for each area was estimated as the average ΔΔ*G* of comprising amino acid residues from three MD simulations. The hydropathy plots for MBP as well as *bovine* rhodopsin (as an example protein with clearly segmented TM and water-soluble regions) were also analyzed using the hydrophobicity scale. The hydrophobic regions of the rhodopsin in the hydropathy plot match those of the protein’s TM domains provided in OPM database (PDB ID 1U19).

### Density functional theory (DFT) calculations

We performed the mixed-reference spin-flip time-dependent DFT (MR-SF-TDDFT) at the BHHLYP/6-31G* level of theory with a conductor-like polarizable continuum model (C-PCM), as implemented in the GAMESS program package^111^. Since a system with an oxygen molecule contains with nearly degenerate π* orbitals which produce doubly degenerate states, the electron correlation effects are crucial for electronic structure calculations, which can be handled by the MR-SF-TDDFT calculation. The excitation energies for singlet and triplet states of amino acid residue-O_2_ pair geometries in each O_2_-trapping subarea were calculated to obtain the absorption spectra. Atom coordinates for the calculations were extracted from the MD trajectories in every 10 ns of the longest dwell-time regions for each area. The smooth absorption spectrum, I(λ), was obtained from I(λ) = (πΓ/2)Σ_i_ ε_osc_^i^ L_i_(λ), where ε_osc_^i^ is the oscillator strength of the i^th^ configuration and L_i_(λ) is the normalized Lorentzian function centered at λ_i,abs_ with the broadening Γ, *i.e.*, L_i_(λ) = 2Γ/(π(4(λ–λ_i,abs_)^2^+Γ^2^)). 40 nm was chosen for the broadening parameter.

### Electron paramagnetic resonance (EPR) spectroscopy

We performed the EPR measurements using 2,2,6,6-tetramethylpiperidine (TEMP), which is commonly used as a ^1^O_2_ spin trapper^112^, at room temperature. The oxidation of TEMP by ^1^O_2_ produces 2,2,6,6-tetramethylpiperidine 1-oxyl (TEMPO) free radical, which exhibits the characteristic EPR signal. 30 mM TEMP in DMSO was mixed with DI water or 50 mM tryptophan in DI water at a 1:3 ratio (v/v) (total volume of 20 µl). The mixtures were incubated for 6 min, with or without exposure to blue light (λ_peak_ = 446 nm, Extended Data Fig. 1 for the full spectrum), and then transferred to EPR capillary tubes. The EPR signals were acquired using EMXplus-9.5/2.7 spectrometer (Bruker) at Kyungpook National University Instrumental Analysis Center in Daegu, Korea. Four scans were conducted for each sample with the following instrument parameters: sweep width, 100 G; power, 0.502 mW; modulation amplitude, 1.0 G; time constant, 81.92 ms; conversion time, 30 ms; sweep time, 61.44 s.

### Monte Carlo (MC) analysis

The atom coordinates of MBP were extracted from a MD trajectory in every 10 ns and then averaged for each atom. The time-averaged coordinates of all side-chain atoms for each residue were further averaged to assign the residue position. The residues from each type were randomly sampled and the distances between the oxidized residues and the sampled residues were averaged over all possible combinations in each sampling. The numbers of sampled residues were one to eight as the total number of the benchmark Trp residues is eight. The sampling number was 10^3^ for each number of sampled residues. All possible cases were analyzed in case that its total number was less than the sampling number. The calculation was iterated to construct a probability density distribution for each residue type. This MC analysis was also conducted for establishing the probability density distributions for arbitrary residues sampled from the entire protein. The sampling number was 10^6^ for each number of sampled residues. Likewise, all possible cases were analyzed in case that its total number is less than the sampling number. The mean value of the average distances analyzed for each residue type was used to calculate the top proximity percentage in Fig. 6d.

## Acknowledgments

This work was supported and funded by the National Research Foundation of Korea (2020R1C1C1003937 to D.M.; 2021R1A2C2009504 to T.-H.K.; 2021R1A3B1077184 to S.K.M.; 2021R1C1C1010943 and 2022R1A4A1033471 to J.-M.C.), POSCO TJ Park Foundation (POSCO Science Fellowship to J.-M.C), National Cancer Center (HA22C010100 to T.-H.K.), and the Institute for Basic Science (IBS-R022-D1 to K.M.).

## Author contributions

D.M. conceived and supervised the project. S.K. and D.M. designed the single-molecule tweezer experiments. S.K., V.W.S., and W.C.B.W. performed the single-molecule tweezer experiments. S.K., V.W.S., and D.M. analyzed the single-molecule experiment data. S.K. and H.J. expressed and purified the protein. S.K., M.P., and C.L. performed the ROS measurements. E.K. and D.M. performed and analyzed the MD simulations. J.-M.C. reviewed the MD simulation results. Y.-G.E. and J.-M.C. performed the free energy calculations. E.K. and D.M. performed the protein cavity analysis. C.U.K. reviewed the protein cavity analysis. S.H.K. and S.K.M. performed the DFT calculations. M.P., G.Y., and T.-H.K. performed the EPR measurements. M.P., B.-G.K., and T.-H.K. prepared the samples for the LC-MS/MS analysis. B.-G.K. and K.M. performed the LC-MS/MS. M.P., E.K., S.K., B.-G.K., T.-H.K., and D.M. analyzed the LC-MS/MS data. E.K. and D.M. performed the MC analysis. S.K., E.K., M.P., S.K.M., T.-H.K., and D.M. prepared the manuscript, with input from S.H.K., B.-G.K., C.U.K., and J.-M.C.

## Data availability

All data that support the findings of this study are available in the main text, Extended Data Figures, and Supplementary Information. The mass spectrometry data have been deposited to the ProteomeXchange Consortium via the PRIDE partner repository with the dataset identifiers PXD047549 and 10.6019/PXD047549. Other raw data have been deposited to figshare (https://doi.org/10.6084/m9.figshare.24751755).

## Code availability

Codes for the analyses based on MD and MC simulations will be available in Code Ocean upon request for peer review.

## Extended Data Figures

**Extended Data Fig. 1.**
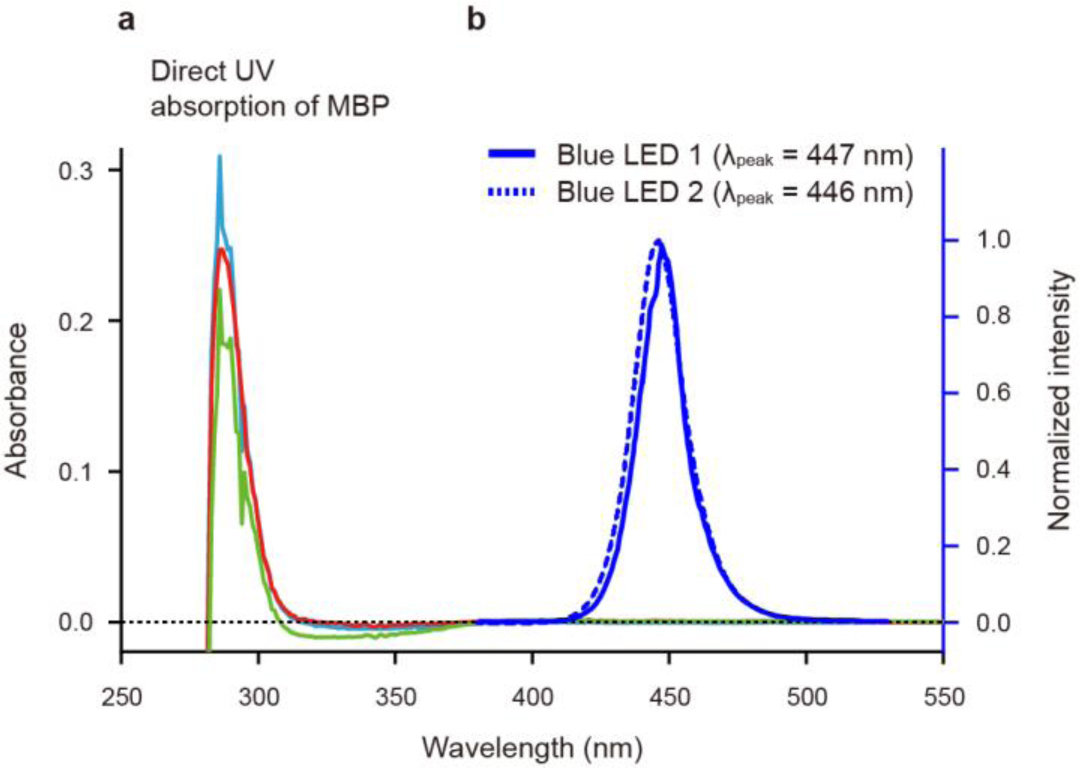
| Absorption spectra of MBP and light spectra used in our study. (a) Absorption spectra of MBP measured using a spectrophotometer. The absorbance of 10 μM MBP in Buffer B (50 mM HEPES, pH 7.6, 100 mM KCl, 5 mM MgCl_2_; used in the foldability measurement) was measured in the range of 250 to 550 nm with a 1-nm interval using the absorbance scan mode of Infinite M200 PRO. The spectra were corrected by subtracting the blank spectra of Buffer B. Three different colors indicate triplicate measurements. (b) Spectra of light sources used in our study. The blue LED 1 (9.2 mW/cm^2^) is used in the single-molecule tweezer experiments, while the blue LED 2 (16 mW/cm^2^) is used in the LC-MS/MS analysis, ABDA assay, and EPR measurements. LED indicates light-emitting diode.

**Extended Data Fig. 2.**
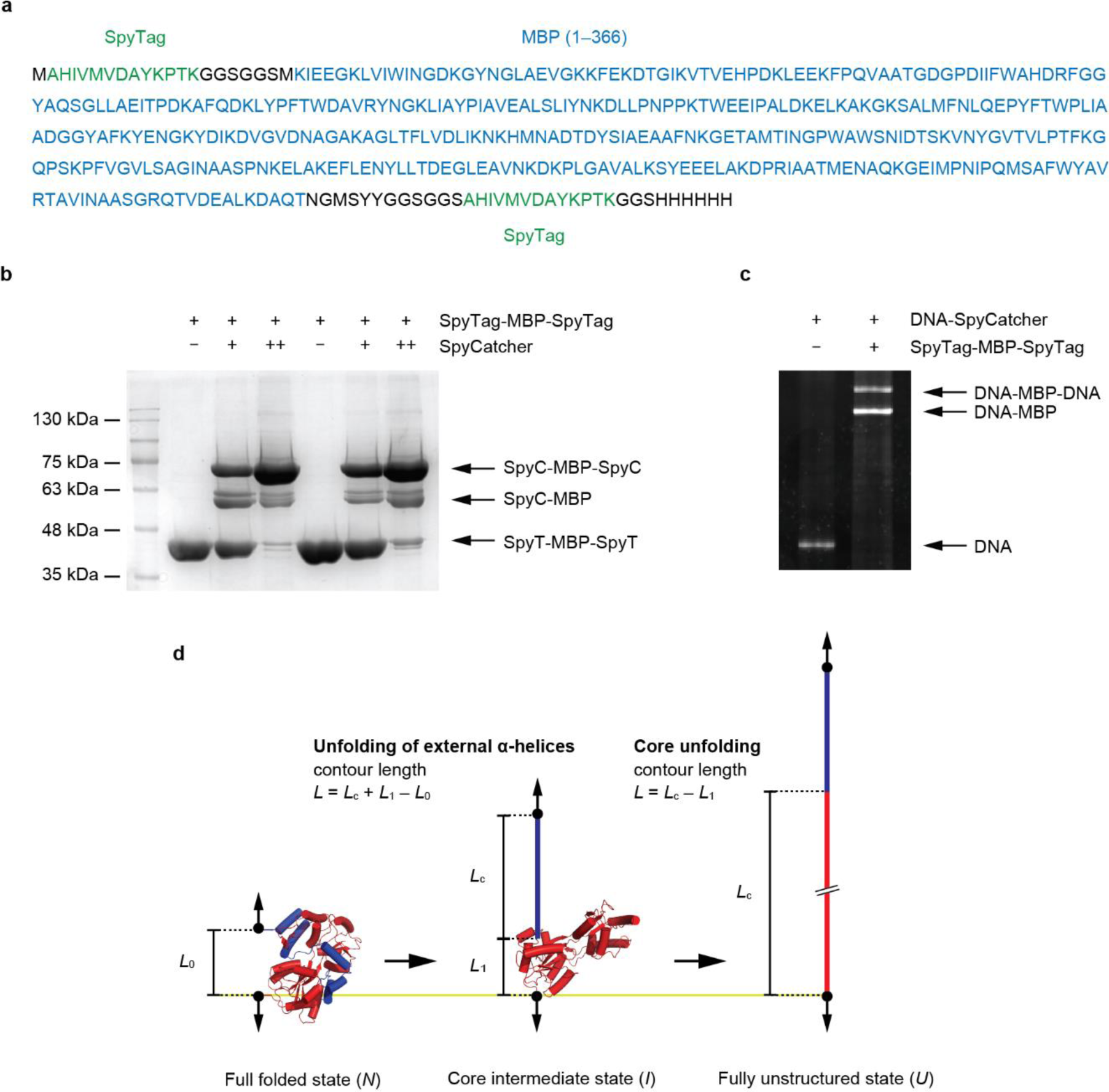
| Sample preparation and MBP unfolding. (a) Amino acid sequence of maltose-binding protein (MBP) construct used in our study. The residue numbers indicate those following the signal peptide sequence. (b) Protein purification and binding test of SpyCatcher (SpyC) to SpyTag (SpyT) (10% SDS-PAGE). The notation “++” indicates a higher concentration of SpyC. (c) DNA handle attachment to MBP. DNA handles conjugated to SpyC are linked to MBP through the binding between SpyC and SpyT. Only the doubly-handled construct with an azide at one end and 2xbiotin at the other end is tethered to the sample chamber surface and magnetic beads (6% SDS-PAGE). (d) Cartoon of MBP unfolding. The unfolding of the external α-helices at the C-terminus (286–366; 81 residues) is followed by the complete unfolding of remaining core structure (7–285; 279 residues). The contour length (*L*) for each state was the contour length of polypeptide portion (*L*_c_) corrected by *L*_0_ and *L*_1_, which are the end-to-end distances of the corresponding protein structures (= 4.21 and 1.62 nm from PDB ID 1ANF).

**Extended Data Fig. 3.**
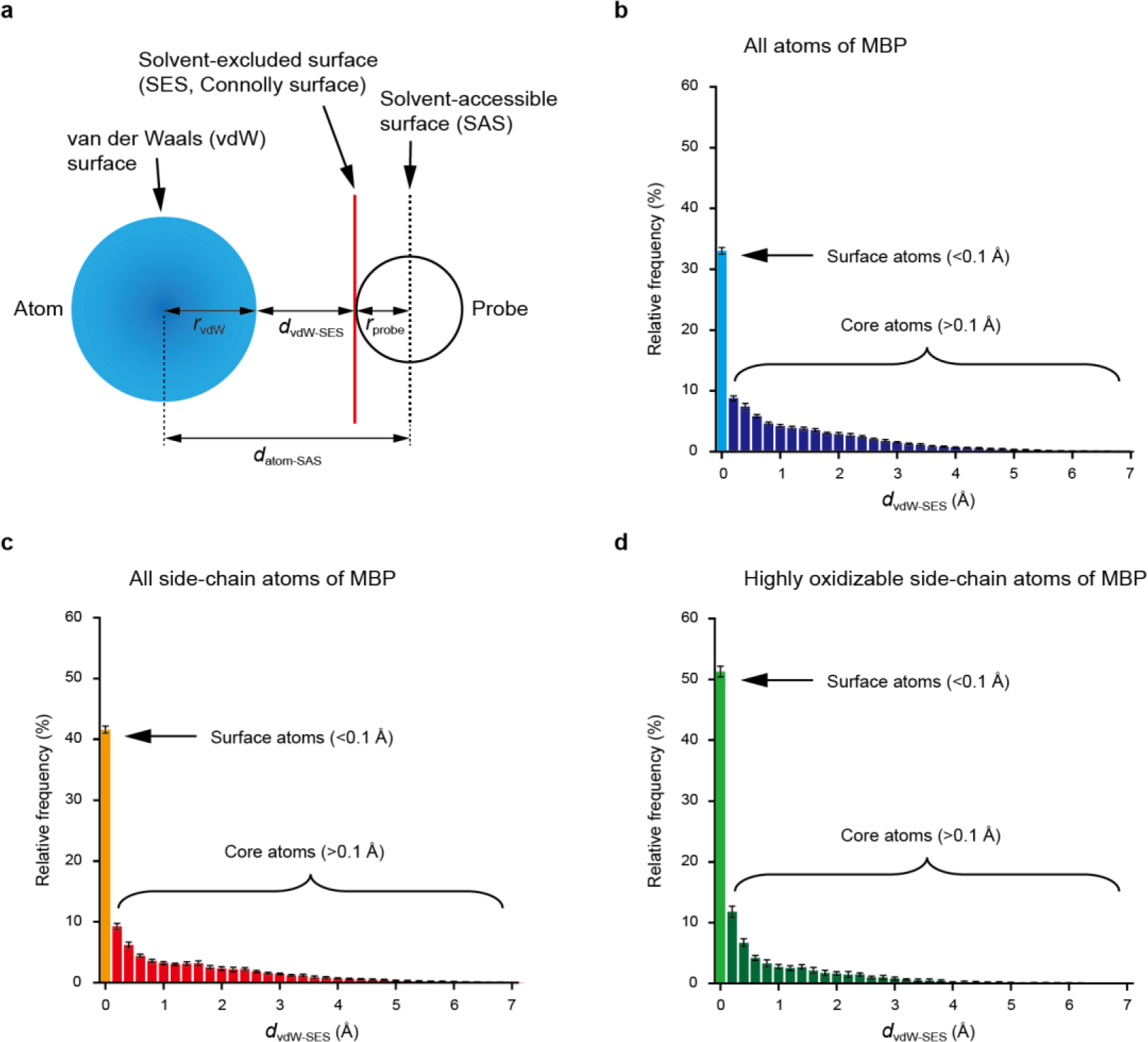
| Estimation of protein surface and core atoms. (a) Illustration depicting protein surfaces and relevant distances using Gromacs solvent-accessible surface area (SASA) module (b–d) Relative frequency of *d*_vdW-SES_ for all atoms of MBP (b), all side-chain atoms of MBP (c), and highly oxidizable side-chain atoms of MBP (d). Refer to the reference in Methods for the highly oxidizable side-chain atoms. For each case, the protein surface or core atoms were classified based on *d*_vdW-SES_ values smaller or larger than 0.1 Å, respectively. The percentages of the surface atoms were estimated as 33.1±0.5%, 41.6±0.6%, and 51.3±0.9% (mean ± SD) for each case, respectively.

**Extended Data Fig. 4.**
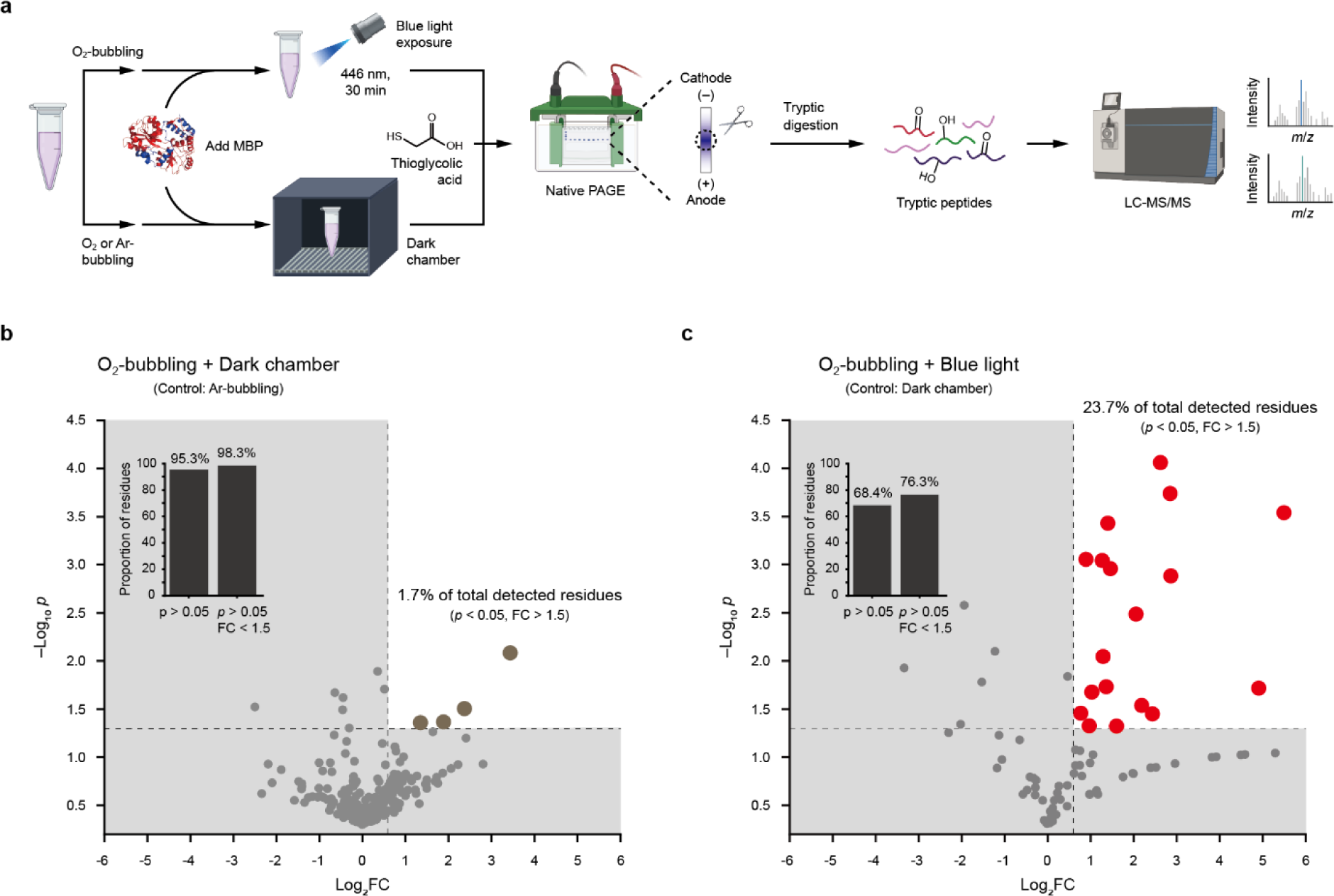
| LC-MS/MS procedure and volcano-plot analysis. (a) Flow diagram of liquid chromatography-tandem mass spectrometry (LC-MS/MS). (b) Volcano plot of oxidized residues under the condition of O_2_-bubbling and incubation in a dark chamber (control: Ar-bubbling condition). The statistically significant oxidized residues (*p*-value (*p*) < 0.05) with a fold change (FC) more than 1.5 correspond to the brown data points in the white background area, accounting for only 1.7% of total detected residues. The residues outside the area or only with *p* > 0.05 account for 98.3% or 95.3% of total detected residues, respectively (inset histogram). (c) Volcano plot of oxidized residues under the condition of O_2_-bubbling and blue light exposure (control: dark chamber condition). The statistically significant oxidized residues (*p* < 0.05) with a FC more than 1.5 correspond to the red data points in the white background area, accounting for 23.7% of total detected residues. The residues outside the area or only with *p* > 0.05 account for 76.3% or 68.4% of total detected residues, respectively (inset histogram).

**Extended Data Fig. 5.**
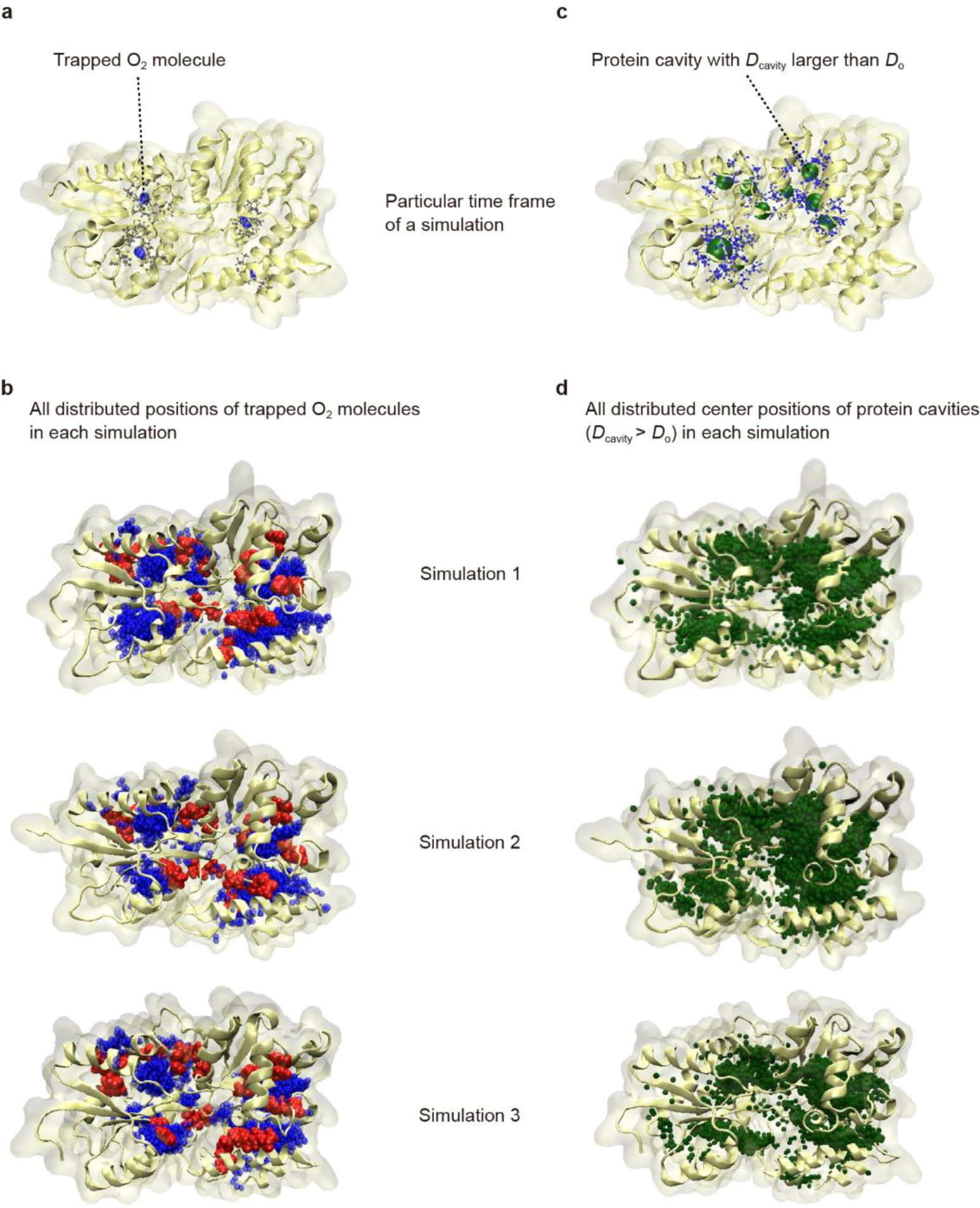
| Structural distribution of trapped O_2_ molecules and protein cavities. (a,b) Trapped O_2_ molecules in a particular time frame of a MD simulation (a) and all distributed positions of trapped O_2_ molecules in each MD simulation (b). (c,d) Protein cavities with *D*_cavity_ larger than *D*_o_ in a particular time frame of a MD simulation (c) and all distributed positions of center positions of the protein cavities in each MD simulation (d). *D*_cavity_ and *D*_o_ represent the cavity diameter and the kinetic diameter of O_2_ (3.46 Å), respectively.

**Extended Data Fig. 6.**
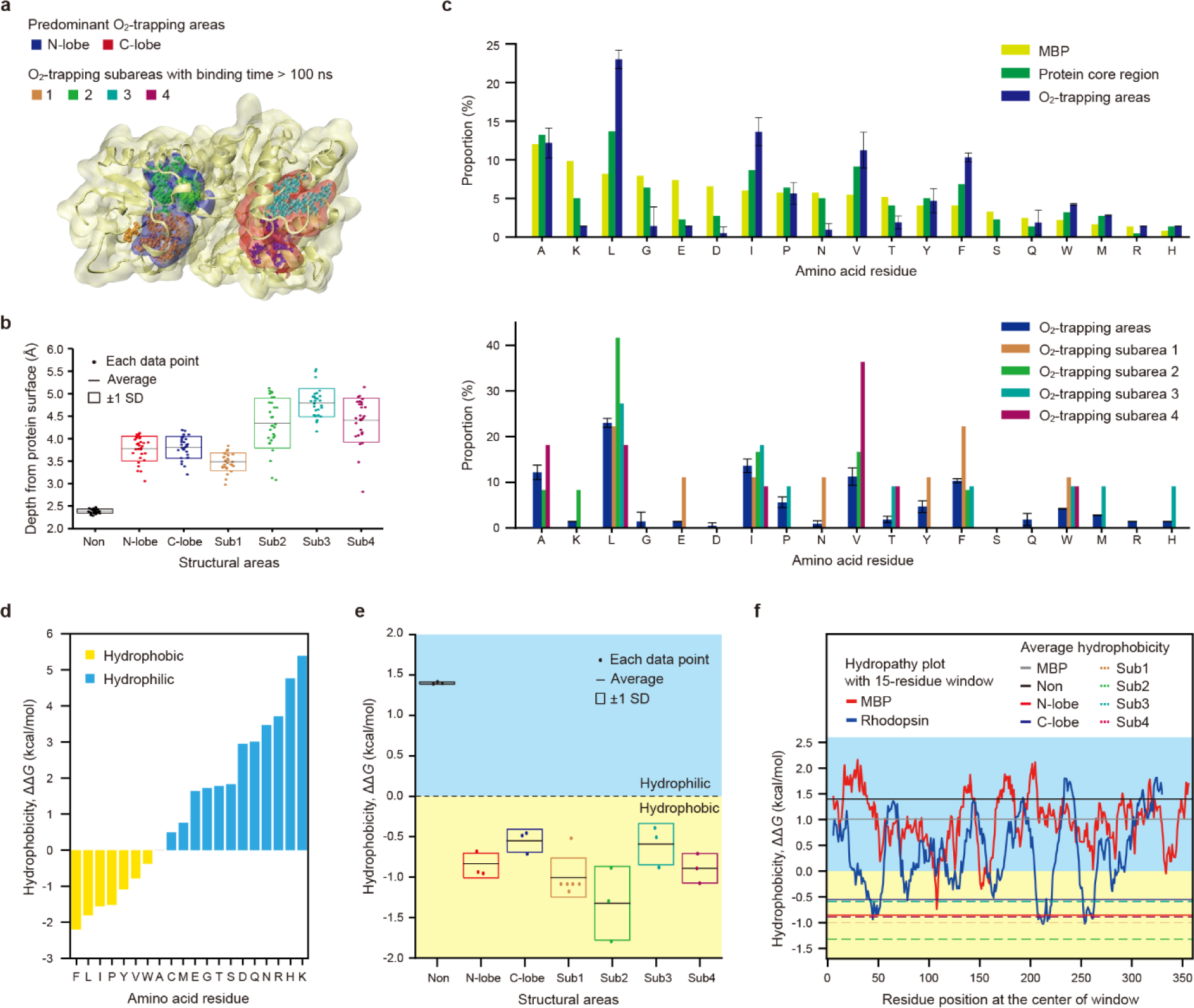
| Depth and hydrophobicity of predominant O_2_-trapping (sub)areas. (a) Predominant O_2_-trapping (sub)areas determined by the procedures described in Supplementary Methods. (b) Depths of O_2_-trapping (sub)areas from the protein surface (solvent-excluded surface). The data were obtained from three MD simulations in every 100 ns (*n* = 30 for each area). (c) Proportion of amino acid residues within specific structural areas. For the O_2_-trapping areas, each bar represents the average of those from three MD simulations (mean ± SD). (d) Moon-Fleming hydrophobicity scale used in hydrophobicity analyses (see Methods for more details). (e) Hydrophobicity values of the O_2_-trapping (sub)areas from three MD simulations (*n* = 3–6 for each area). N-/C-lobe and Sub# indicate the O_2_-trapping area in the N- or C-terminal lobe and each O_2_-trapping subarea, respectively. Non indicates the non-binding area which include residues not involved in the predominant O_2_-trapping areas. (f) Hydropathy plots for MBP and *bovine* rhodopsin. Hydrophobicity values were averaged with a moving 15-residue window. In the analyses shown here, the maximum residue set from three MD simulations was used for each O_2_-trapping subarea (Supplementary Table 2).

**Extended Data Fig. 7.**
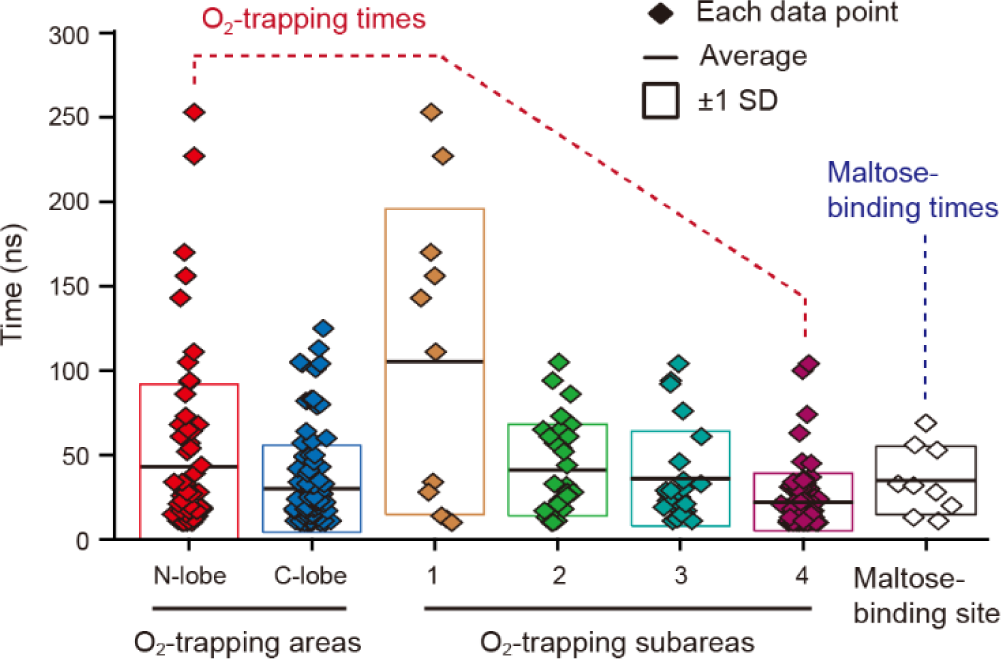
| Distribution of O_2_-trapping times and maltose-binding times. The dwell times of trapped O_2_ molecules within each structural area were obtained through the procedure shown in Supplementary Figs. 6–8. The maltose-binding times were obtained through the procedure shown in Supplementary Fig. 9. Refer to Supplementary Methods for more details.

**Extended Data Fig. 8.**
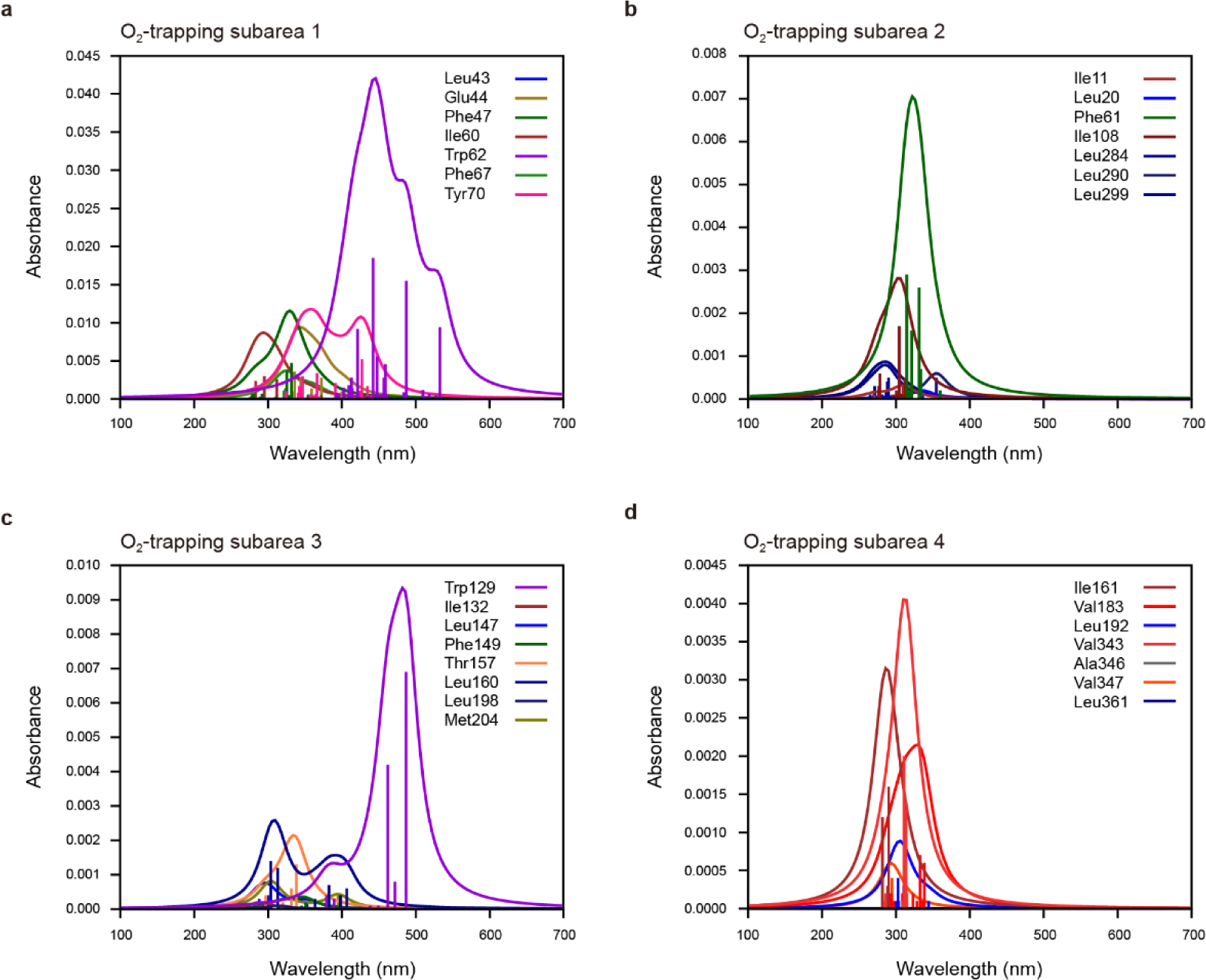
| Absorption spectra of residue-O_2_ pairs in each O_2_-trapping subarea. The Trp62-O_2_ pair in subarea 1 shows the highest absorption at 445 nm, among all residue-O_2_ pairs. The Trp129-O_2_ pair in subarea 3 and the Phe61-O_2_ pair in subarea 2 show relatively high absorptions at 480 nm and 320 nm, respectively, compared to other residues in each subarea. Only the absorption peaks for the Trp62-O_2_ and Trp129-O_2_ pairs closely match the illumination wavelength in the single-molecule tweezer experiments (λ_peak_ = 447 nm; Extended Data Fig. 1 for the full spectrum). The amino acid residues are denoted as 3-letter code.

**Extended Data Fig. 9.**
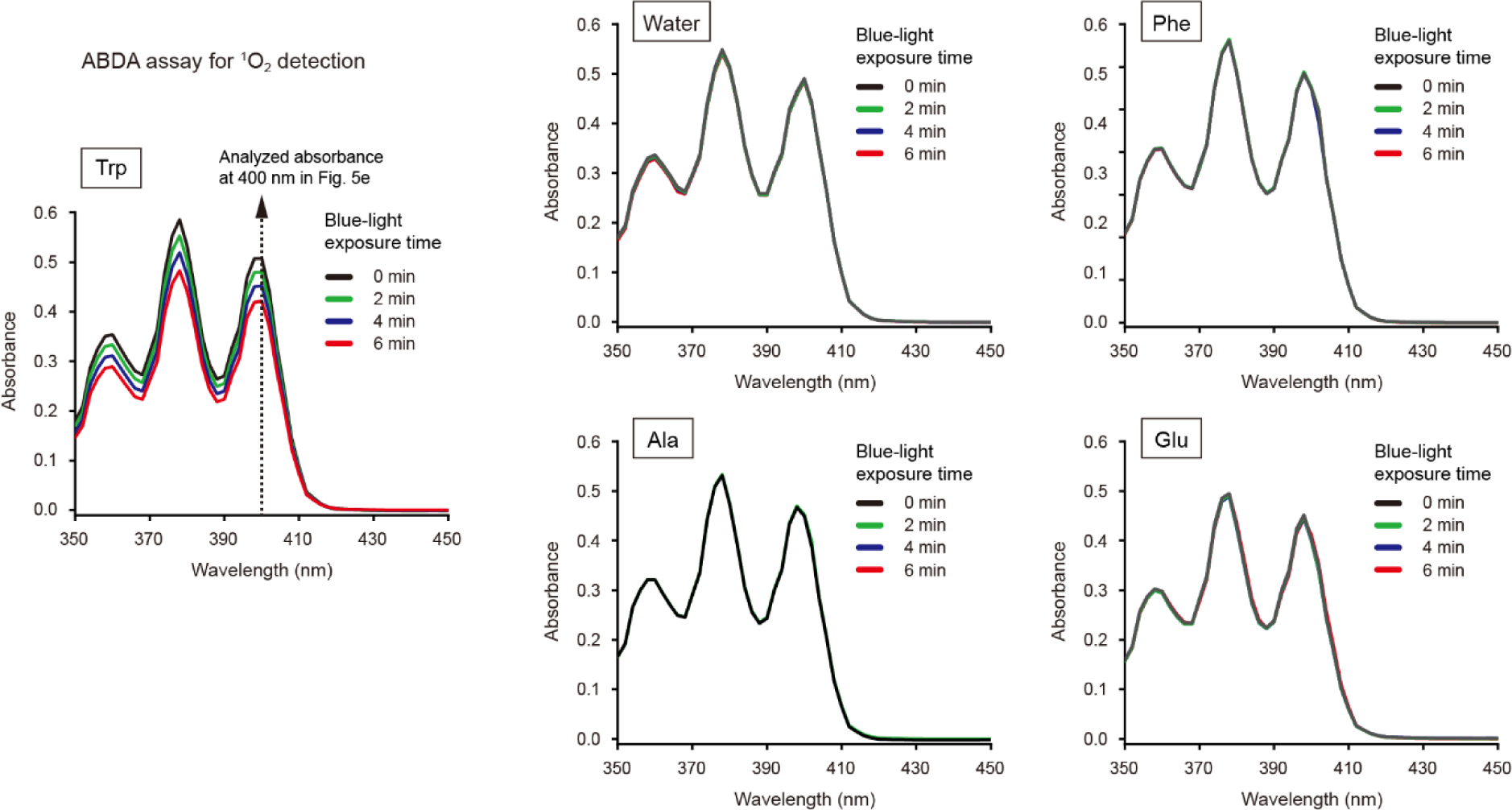
| ABDA assay for free amino acids under blue light. 25 mM of each amino acid in deionized (DI) water (or only DI water) and 10 mM ABDA in DMSO were mixed at a ratio of 99:1 (v/v), and then exposed to blue light (λ_peak_ = 446 nm; Extended Data Fig. 1 for the full spectrum) for 0, 2, 4, and 6 min. The absorbance was measured in the range of 350 to 450 nm with a 2-nm interval using the absorbance scan mode of Infinite M200 PRO. The absorbance values were corrected by that of 1% (v/v) DMSO as the blank absorbance. The measurement for each condition was conducted in triplicate and the data were averaged at each wavelength. The oxidation of ABDA by ^1^O_2_ is shown as the absorbance decrease. The amino acids are denoted as 3-letter code.

**Extended Data Fig. 10.**
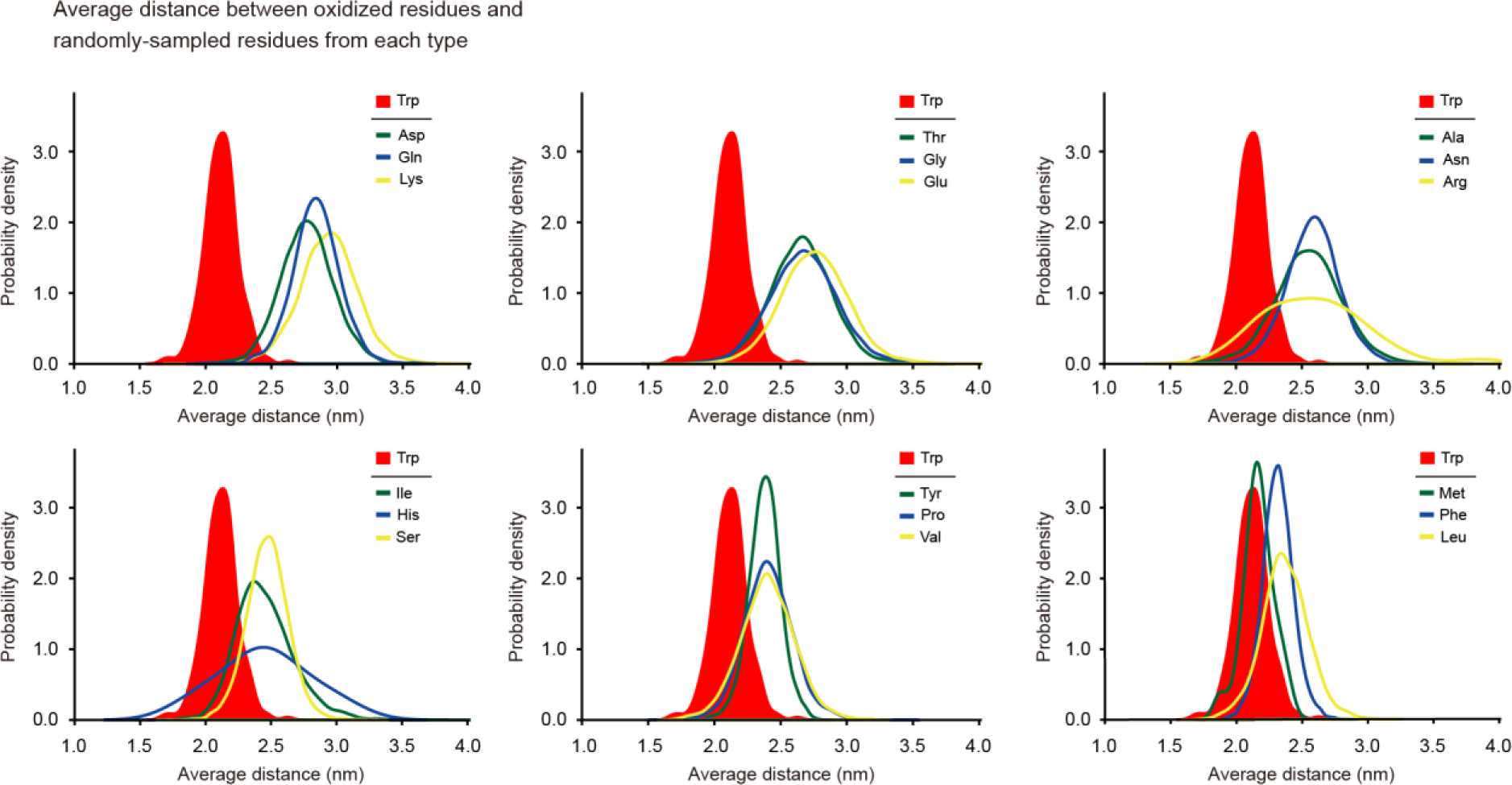
| Proximity between oxidized residues and each residue type. Each plot represents the probability density distribution of average distances between the oxidized residues and the randomly-sampled residues from each residue type. The numbers of sampled residues are one to eight as the total number of the benchmark Trp residues is eight. The sampling number is 10^3^ for each number of sampled residues. All possible cases are analyzed in case that its total number is less than the sampling number. Cys is excluded in this analysis because MBP does not have Cys residues. The amino acid residues are denoted as 3-letter code. Refer to Methods for more details.

