## Supplementary Information for "Hidden route of protein damage through confined oxygen gas"

### Contents


**Supplementary Methods**

**Supplementary Figures**

Supplementary Fig. 1 | Measurement of dissolved $O_2$ concentration and pH

Supplementary Fig. 2 | Distribution of protein cavity diameters

Supplementary Fig. 3 | Determination of predominant $O_2$-trapping areas

Supplementary Fig. 4 | Determination of $O_2$-trapping subareas

Supplementary Fig. 5 | Free energies for $O_2$-trapping events

Supplementary Fig. 6 | Extraction of $O_2$-trapping times for a specific structural area

Supplementary Fig. 7 | Barcode plot generation for $O_2$-trapping areas in each lobe

Supplementary Fig. 8 | Barcode plot generation for $O_2$-trapping subareas

Supplementary Fig. 9 | Analysis of maltose-binding times

**Supplementary Tables**

Supplementary Table 1 | Residues comprising predominant $O_2$-trapping areas

Supplementary Table 2 | Residues comprising $O_2$-trapping subareas

**Supplementary Videos**

Supplementary Video 1 | Representative $O_2$-trapping events

Supplementary Video 2 | Representative maltose-binding events

**References**

### Supplementary Methods

**Determination of predominant $O_2$-trapping (sub)areas**

To determine the predominant $O_2$-trapping areas (Supplementary Fig. 3 and Supplementary Table 1), the minimum distance between atoms of MBP and $O_2$ molecules was first extracted in every 1 ns from a MD simulation. We collected all time points, during which the distance remains less than a cutoff distance for longer than a cut-off time span. 3.5 Å was used as the cutoff distance to include only the close binding. The cutoff distance value of 3.5 Å has been used to determine the atoms involved in protein-ligand interactions[1-3]. 10 ns was used as the cutoff time span to exclude a large number of transient interactions and only consider more sustained binding events. All protein residues of which the atoms are less than the cutoff distance from $O_2$ atoms were extracted from the binding time points in every 1 ns. From a histogram of total counted times for the extracted residues, the top 32.9±1.7% (mean ± SD; $n$ = 3 simulations) of the residues were selected for the predominant $O_2$-binding residues, covering 90% of the total counted times of all extracted residues. The selected residues can be considered as the residues that bind to $O_2$ as predominantly as 90% of the total binding times. The $O_2$-binding residues comprise the $O_2$-trapping areas in the N- and C-terminal lobes of MBP. We conducted this analysis for three MD simulations and determined the common $O_2$-trapping areas. To determine the $O_2$-trapping subareas that more tightly bind to $O_2$ (Supplementary Fig. 4 and Supplementary Table 2), only the binding time regions longer than 100 ns were analyzed (7.6% of all binding time regions). All protein residues of which the atoms are less than the cutoff distance from $O_2$ atoms were extracted from the binding time points in every 1 ns. The residues tightly binding to $O_2$ were selected from the highest counted residue by the cutoff number of residues, from a histogram of total counted time for each extracted residue. The cutoff number was defined as the maximum number of residues binding to $O_2$ at a certain time point. From three MD simulations, total 15 cases with >100-ns binding times were analyzed and classified into four distinct $O_2$-trapping subareas based on residue overlap.

**Free energy calculations for $O_2$-trapping events**

We conducted the free energy calculations using the alchemical transformation method applied to the generated MD trajectories. The initial structures were sampled from the MD trajectories, where $O_2$ is trapped within the $O_2$-trapping subareas for the longest dwell times (Supplementary Table 2). For the alchemical transformation, a dummy $O_2$ molecule was introduced into the surrounding solution. The simulations employed 20 windows with different λ values. Over the range between $\lambda=0$ and $\lambda=1$, the interactions of the trapped $O_2$ molecule within the complex were decreased, while those of the dummy $O_2$ were recovered. Within each window, the systems were equilibrated in the following sequence: (1) energy minimization, utilizing the steepest descent algorithm for a maximum of 5000 steps; (2) 100-ps NVT equilibration, using Langevin dynamics[4] with a reference temperature of 300K; (3) 100-ps NPT equilibration, employing the Berendsen algorithm[5]. Throughout the two equilibration steps, position restraints were applied to the trapped $O_2$ molecule. A production run of 10 ns was conducted using the Parrinello-Rahman pressure coupling scheme[6]. The multiple Bennett acceptance ratio (MBAR) method[7,8] was employed to estimate the free energies accompanied with the alchemical transformations (Supplementary Fig. 5).

**Analysis for $O_2$-trapping times and maltose-binding times**

Representative MD trajectories and barcode plots for the $O_2$-trapping (sub)areas were obtained through the same procedure used to determine the predominant $O_2$-trapping areas, but only with residues of each structural region considered (Supplementary Figs. 6–8). The $O_2$-trapping times for each structural region were extracted from the barcode plots from three 1-μs MD simulations. The maltose-binding times were obtained only for the specific binding to the maltose-binding site (Supplementary Fig. 9). The comprising residues of the maltose-binding site (D14, K15, W62, D65, R66, E111, E153, Y155, W230, W340) were selected from previous structural studies[9,10]. The structure of maltose (45 atoms) is relatively large and complex compared to $O_2$, so the binding criteria based on the minimum distance between atoms can be misleading in determining the specific/fit binding to the binding site. Instead, the distance between the center of mass of the selected residues and maltose was utilized for the binding criteria[11] and calculated at every 1 ns from a 15-μs MD simulation. The corresponding barcode plot was generated for the time regions during which the distance remains less than 8 Å for longer than 10 ns. The 8-Å cutoff distance can distinguish the specific and nonspecific bindings (Supplementary Fig. 9c). Transient interactions shorter than 10 ns were excluded, same as in the analysis of $O_2$-trapping times. Representative maltose-binding events are visualized in Supplementary Video 4.

### Supplementary Figures

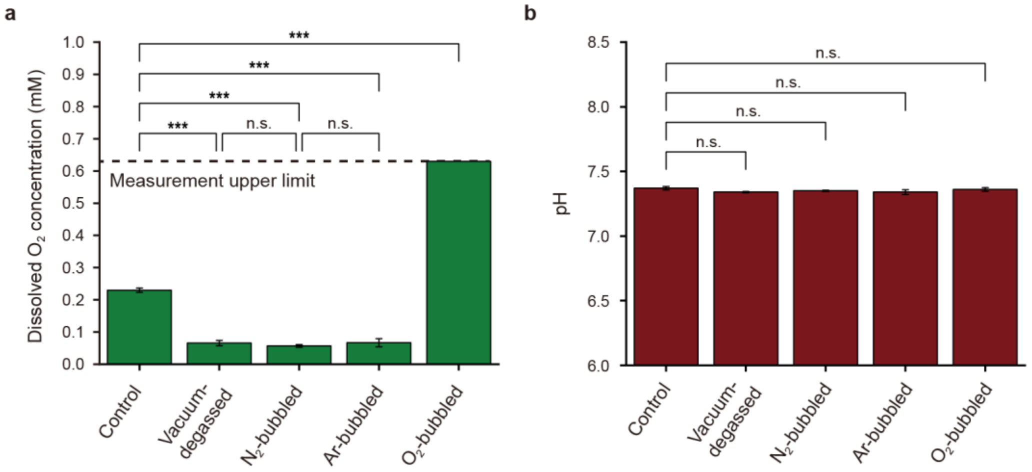


**Supplementary Fig. 1 | Measurement of dissolved $O_2$ concentration and pH.** (a) Measurement of dissolved $O_2$ (DO) concentrations. The control buffer was 50 mM HEPES, pH 7.4, 100 mM KCl, 5 mM $MgCl_2$ used in the foldability measurement. The DO concentration in the control buffer was measured to be 0.23±0.007 mM (mean ± SD, $n$ = 9). After vacuum degassing or gas bubbling, the DO concentrations of the buffers were immediately measured ($n$ = 3–6). The DO concentration of the $O_2$-bubbled buffer corresponds to the measurement upper limit (dashed line). (b) Measurement of pH. After vacuum degassing or gas bubbling, pH of each buffer was immediately measured ($n$ = 3 for each). The statistical test indicates that the pH did not significantly change after the vacuum degassing or gas bubbling. One-way ANOVA with post-hoc Tukey test (*** for $p < 0.001$, n.s. for $p > 0.05$).

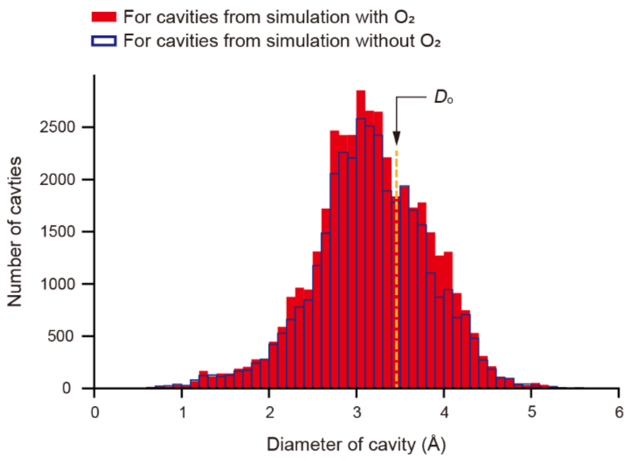


**Supplementary Fig. 2 | Distribution of protein cavity diameters.** The diameters and center positions of protein cavities were obtained using Voronoia**.** Refer to Methods for more details. $D_0$ indicates the kinetic diameter of $O_2$ ($D_0$ = 3.46 Å).

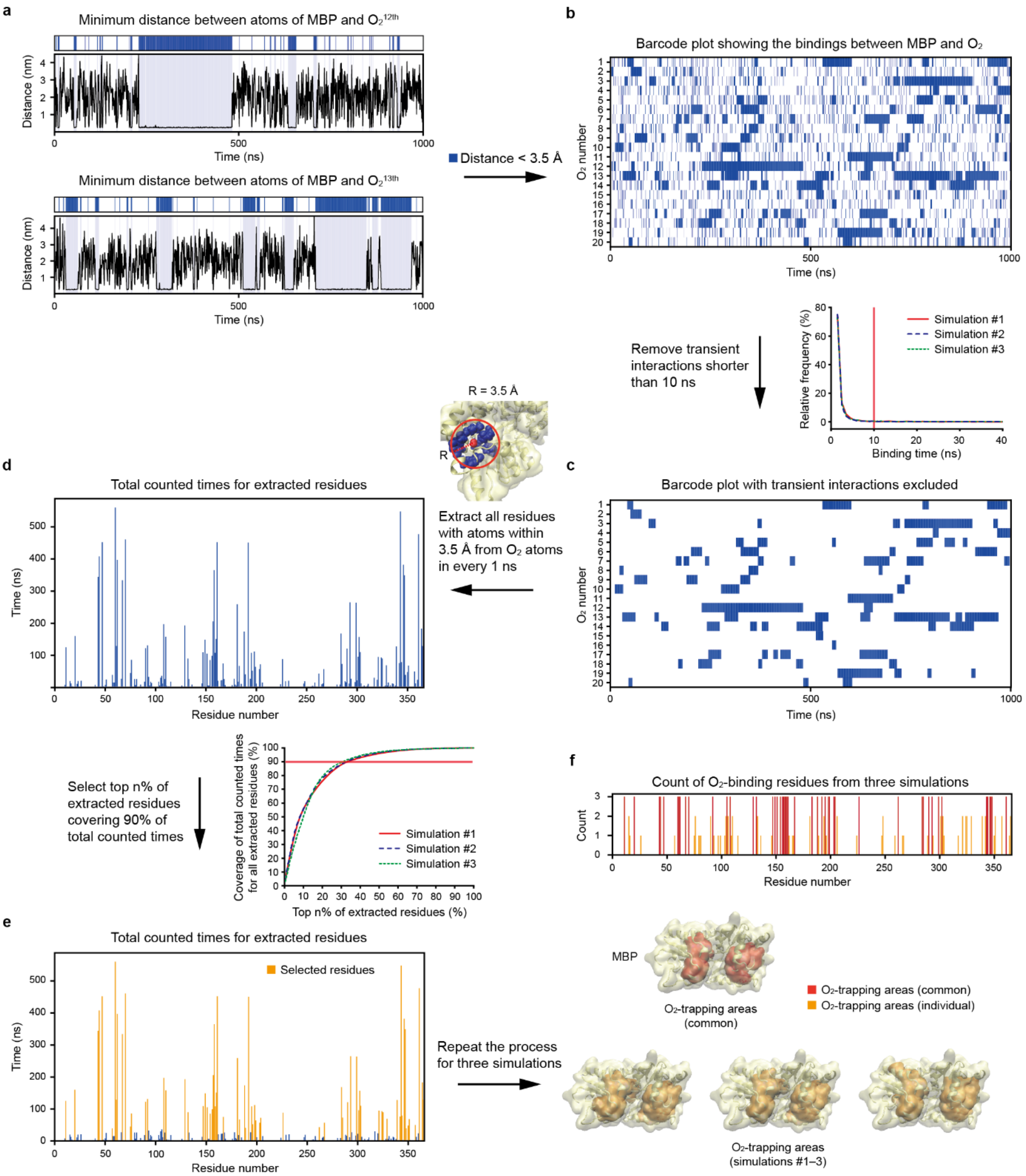


**Supplementary Fig. 3 | Determination of predominant $O_2$-trapping areas.** Refer to Supplementary Methods. (a) Representative time traces showing the minimum distances between atoms of maltose-binding protein (MBP) and $O_2$. The corresponding barcode plots above the traces indicate the binding times during which the distances are less than 3.5 Å. (b) Barcode plot illustrating the bindings between MBP and all $O_2$ molecules in a 1-μs MD simulation. Each row of the barcode plot was obtained from the plots in panel a. The inset in the lower right corner displays the relative frequency as a function of binding time. The relative frequency rapidly decays, with the binding times shorter than 10 ns accounting for 96.6±0.5% (mean ± SD; $n$ = 3 simulations). (c) Barcode plot with transient interactions excluded. Binding times shorter than 10 ns were removed in the barcode plot. (d) Histogram showing the total counted times for $O_2$-binding residues. All protein residues with atoms within 3.5 Å from $O_2$ atoms were extracted for all binding-time points in every 1 ns, and then counted for the 1-μs simulation time span. (e) Determination of predominant $O_2$-binding residues. The top 32.9±1.7% (mean ± SD; $n = 3$ simulations) of the extracted residues were selected for the predominant $O_2$-binding residues (denoted in yellow), covering 90% of the total counted times of all extracted residues (inset in the upper right corner). (f) Predominant $O_2$-trapping areas of MBP. The structural areas for the $O_2$-binding residues are denoted in dark yellow (Supplementary Table 1 for detailed residue information). The common $O_2$-trapping area determined from three simulations are denoted in red. The upper histogram shows the total count for each $O_2$-binding residue in the three simulations.

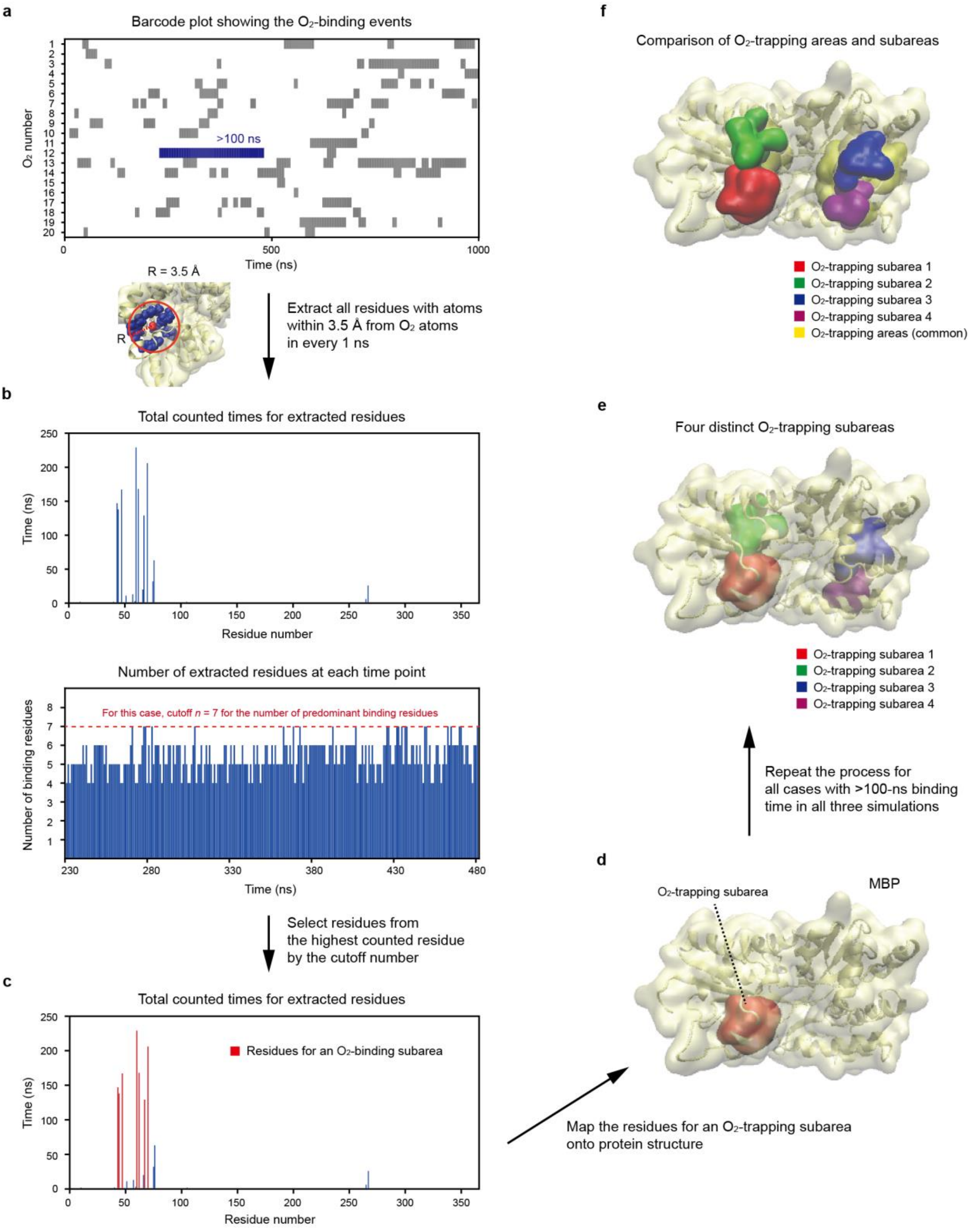


**Supplementary Fig. 4 | Determination of $O_2$-trapping subareas.** Refer to Supplementary Methods. (a) Barcode plot highlighting a binding-time region longer than 100 ns, generated using the procedure shown in Supplementary Fig. 3. (b) Extraction of $O_2$-binding residues during the relatively long binding time. The upper histogram shows the total counted times for the $O_2$-binding residues throughout the binding time. All protein residues with atoms within 3.5 Å from $O_2$ atoms were extracted for the binding-time points in every 1 ns, and then counted for the binding time span. The lower histogram displays the number of extracted $O_2$-binding residues at each time point. The maximum number was set as the cutoff number for the selection of predominant $O_2$-binding residues ($n = 7$ for this example case). (c) Determination of predominant $O_2$-binding residues. Residues predominantly binding to $O_2$ were selected in descending order from the highest counted residue by the cutoff number of residues. (d) An $O_2$-trapping subarea found from the aforementioned procedure. (e) All $O_2$-trapping subareas identified from the analysis for all binding-time regions longer than 100 ns. The $O_2$-trapping subareas were classified into four distinct areas based on residue overlap (Supplementary Table 2). The illustrated areas represent those of the longest binding-time case for each area. (f) Comparison of the predominant $O_2$-trapping areas and subareas. The $O_2$-trapping subareas are overlapped with the broader $O_2$-trapping areas.

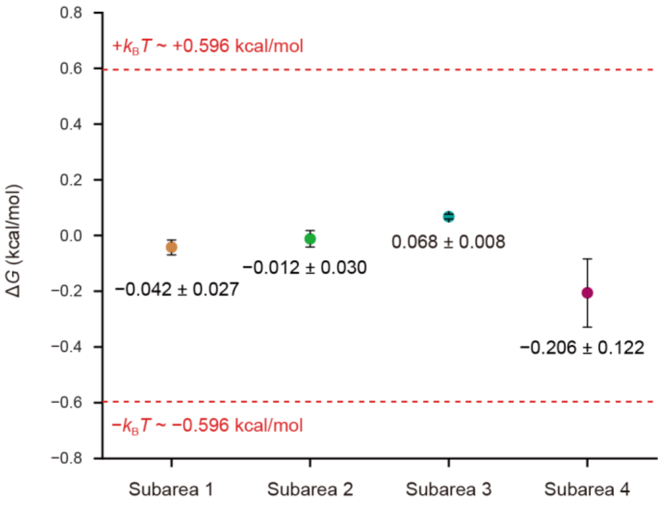


**Supplementary Fig. 5 | Free energies for $O_2$-trapping events.** The $\Delta G$ values were obtained using the alchemical transformation method applied to the MD trajectories, where $O_2$ is trapped within the $O_2$-trapping subareas for the longest dwell times (Supplementary Table 2). Refer to Supplementary Methods for more details. Here, $\Delta G$ is defined as $\Delta G = G_{\text{bound}} - G_{\text{unbound}}$. The estimated $\Delta G$ values for $O_2$-trapping events range from –0.206 to 0.068 kcal/mol. These $\Delta G$ values are below the thermal energy level of ±0.596 kcal/mol, confirming the thermodynamic feasibility of the $O_2$-trapping events.

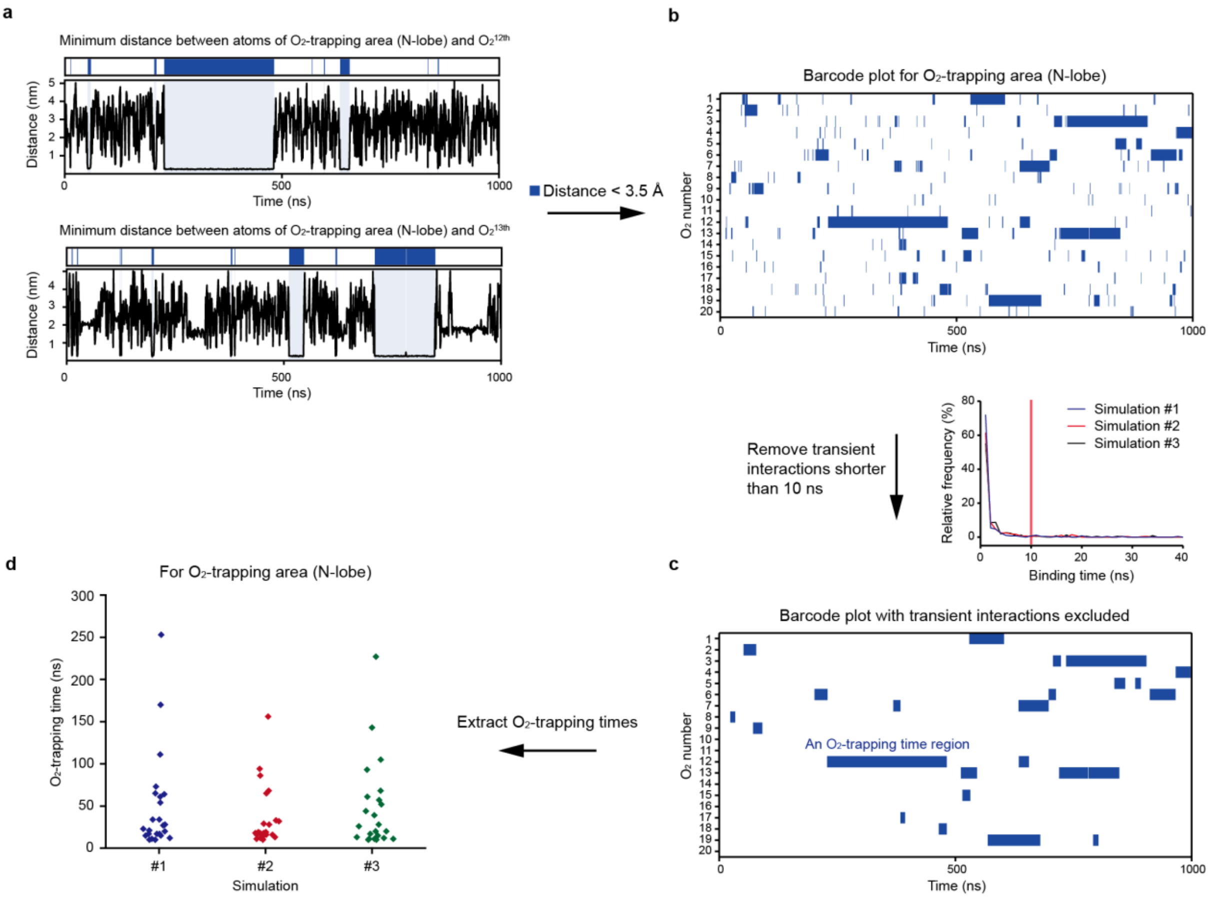


**Supplementary Fig. 6 | Extraction of $O_2$-trapping times for a specific structural area.** Refer to Supplementary Methods. This figure displays an example procedure for the analysis of O-trapping times within a structural region of interest. (a) Representative time traces showing the minimum distances between atoms of the predominant $O_2$-trapping area (N-lobe) and $O_2$ molecules. The corresponding barcode plots above the traces indicate the $O_2$-binding times during which the minimum distances are less than 3.5 Å. Additional traces are available in Supplementary Fig. 7. (b) Barcode plot illustrating the $O_2$-binding times for all $O_2$ molecules. Each row of the barcode plot was obtained from the plots in panel a. The inset in the lower right corner shows the relative frequency as a function of binding time, which rapidly decays within 10 ns. (c) Barcode plot with transient interactions excluded. Binding times shorter than 10 ns were removed in the barcode plot. (d) Distribution of $O_2$-trapping times for the $O_2$-trapping area (N-lobe), obtained from three 1-μs MD simulations. For other $O_2$-trapping (sub)areas, the $O_2$-trapping times were extracted from the respective barcode plots shown in Supplementary Figs. 7,8.

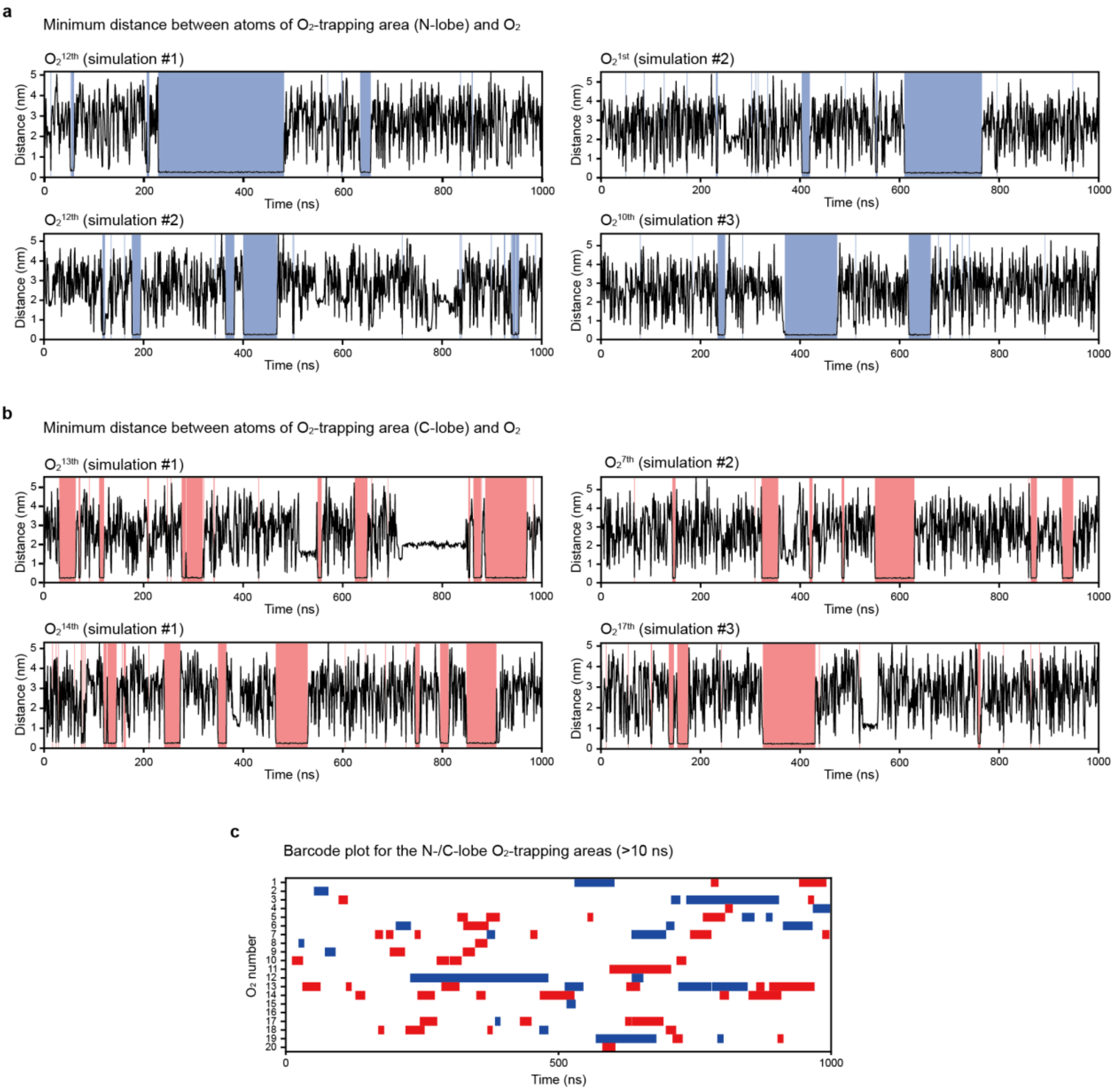


**Supplementary Fig. 7 | Barcode plot generation for $O_2$-trapping areas in each lobe.** (a,b) Time traces of the minimum distance between atoms of the predominant $O_2$-trapping areas and $O_2$. The time spans during which the distance is less than 3.5 Å are colored blue for the N-lobe area (a) and red for the C-lobe area (b). These traces were obtained from three 1-µs MD simulations. (c) Barcode plot for the N-/C lobe $O_2$-trapping areas (illustrative case for simulation #1). The procedure for obtaining the barcode plot is same as shown in Supplementary Fig. 3a–c. All $O_2$-trapping times extracted from the barcode plots of three 1-µs MD simulations are presented in Extended Data Fig. 7.

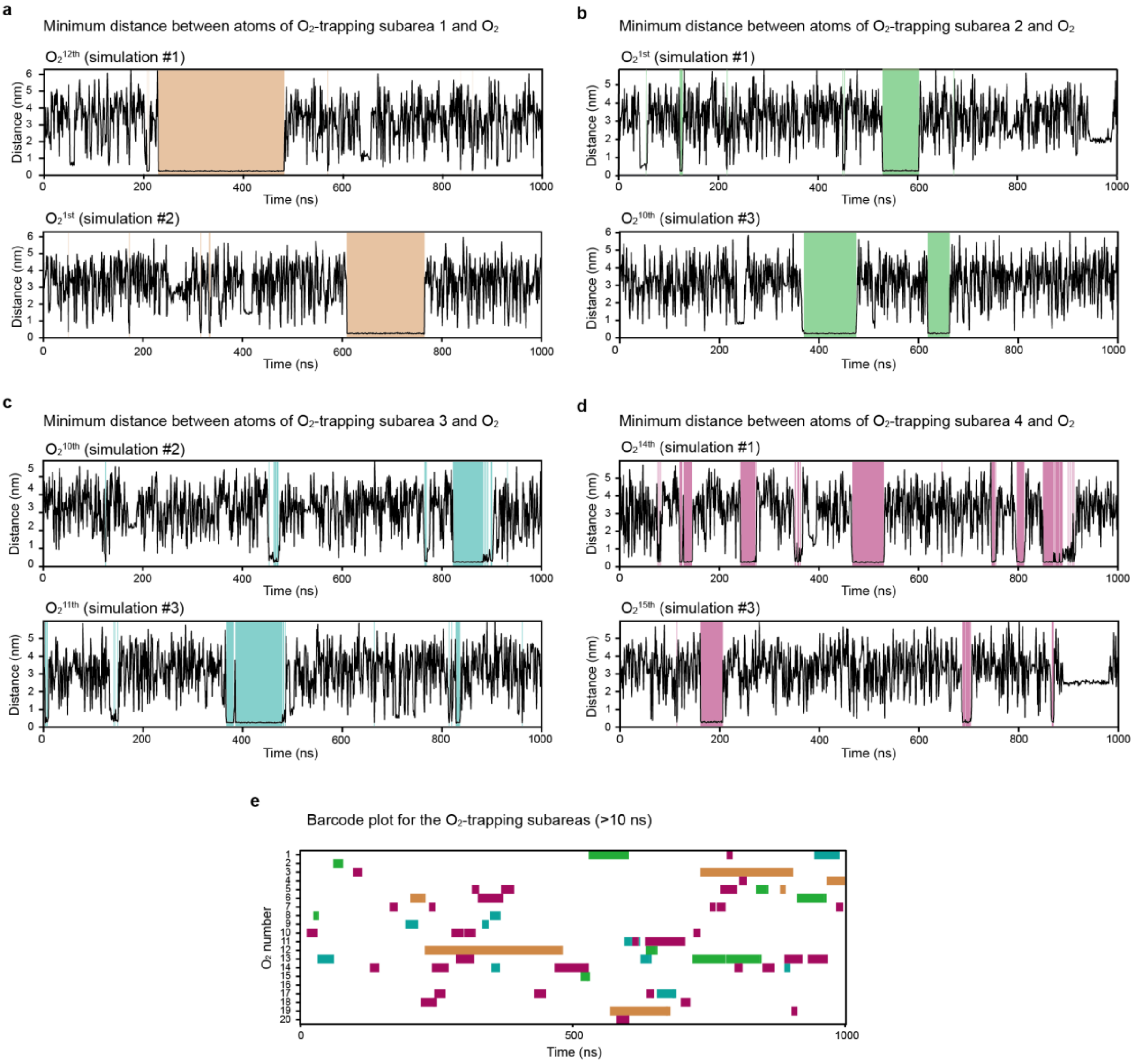


**Supplementary Fig. 8 | Barcode plot generation for $O_2$-trapping subareas.** (a–d) Time traces of the minimum distance between atoms of each $O_2$-trapping subarea and $O_2$. The time spans during which the distance is less than 3.5 Å are colored orange for the subarea 1 (a), green for the subarea 2 (b), cyan for the subarea 3 (c), and purple for the subarea 4 (d). These traces were obtained from three 1-μs MD simulations. The maximum residue set for each $O_2$-trapping subarea was used to plot the traces (Supplementary Table 2). (e) Barcode plot for the $O_2$-trapping subareas (illustrative case for simulation #1). The procedure for obtaining the barcode plot is same as shown in Supplementary Fig. 3a–c. All $O_2$-trapping times extracted from the barcode plots of three 1-μs MD simulations are presented in Extended Data Fig. 7.

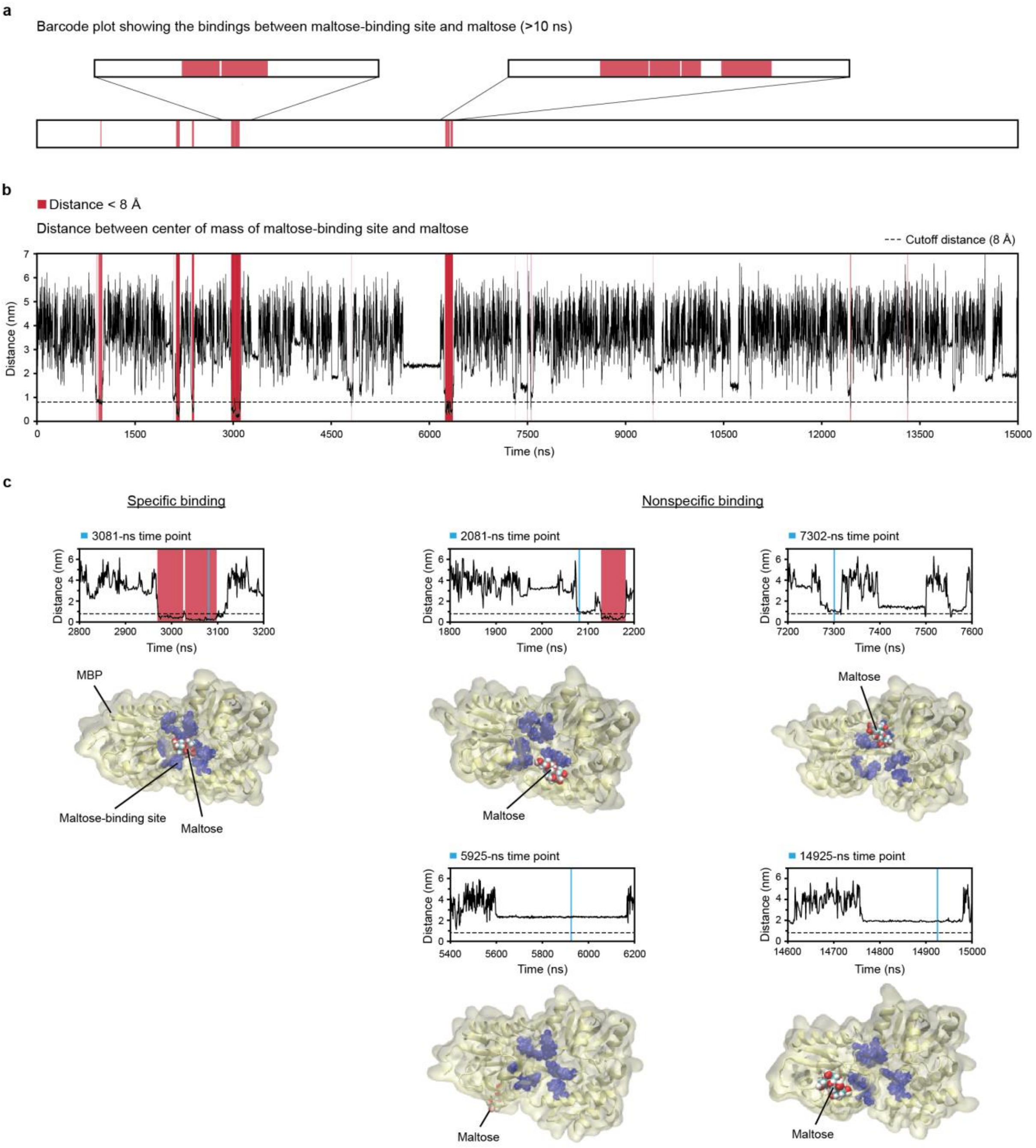


**Supplementary Fig. 9 | Analysis of maltose-binding times.** (a) Barcode plot illustrating the bindings between the maltose-binding site of MBP and maltose. The residues of the maltose-binding site selected for this analysis were D14, K15, W62, D65, R66, E111, E153, Y155, W230, and W340 (see Methods). Transient interactions shorter than 10 ns were removed, same as in the analysis of $O_2$-trapping times. The zoomed-in regions of the binding times are shown above the plot. This barcode plot was generated from the distance plot in panel b. (b) Time trace of the distance between center of mass of maltose-binding site and maltose. The distance was calculated at every 1 ns of a 15-µs MD simulation. The maltose intimately binds to the maltose-binding site during the red-colored time spans with a distance less than 8 Å, as shown in panel c. (c) Snapshots at different time points. The specific binding to the maltose-binding site is observed only in the selected red-colored regions.

### Supplementary Tables

| Simulation # | Residue # |
|---|---|
| 1 | 11, 20, 43, 44, 47, 57, 60, 61, 62, 67, 70, 75, 76, 90, 92, 106, 108, 110, 129, 132, 147, 149, 151, 154, 156, 157, 158, 159, 160, 161, 167, 181, 183, 188, 189, 192, 195, 196, 198, 199, 203, 204, 226, 262, 267, 284, 285, 290, 293, 299, 301, 302, 303, 321, 324, 325, 329, 339, 342, 343, 344, 346, 347, 348, 350, 351, 357, 361, 364, 365 |
| 2 | 11, 15, 20, 43, 44, 47, 60, 61, 62, 67, 70, 76, 90, 92, 97, 99, 104, 105, 108, 109, 110, 129, 132, 133, 147, 149, 151, 154, 156, 157, 158, 159, 160, 161, 167, 181, 183, 188, 192, 194, 195, 198, 199, 203, 204, 206, 226, 248, 262, 284, 285, 290, 293, 294, 299, 301, 302, 303, 304, 321, 324, 325, 339, 342, 343, 344, 346, 347, 348, 350, 357, 361, 365 |
| 3 | 11, 20, 23, 26, 43, 44, 47, 60, 61, 62, 67, 70, 79, 80, 81, 92, 103, 104, 105, 106, 108, 115, 129, 132, 133, 136, 146, 147, 149, 151, 154, 156, 157, 158, 159, 160, 161, 162, 165, 166, 183, 188, 192, 194, 195, 198, 199, 203, 204, 206, 224, 226, 247, 248, 262, 266, 284, 285, 289, 290, 293, 299, 302, 343, 344, 346, 347, 348, 351, 361 |

**Supplementary Table 1 | Residues comprising predominant $O_2$-trapping areas.**

| Area # | Simulation # | Gas # | Residue # | $O_2$-trapping time (ns) |
|---|---|---|---|---|
| 1 | 1 | 3 | L43, E44, F47, I60, W62, F67, Y70 | 169 |
| | | 12 | L43, E44, F47, I60, W62, F67, Y70 | 252 |
| | | 19 | L43, E44, F47, I60, W62, F67, Y70 | 111 |
| | 2 | 1 | L43, E44, F47, I60, W62, F67, Y70, L76 | 155 |
| | 3 | 1 | L43, E44, F47, I60, W62, F67, Y70, N267 | 142 |
| | | 3 | L43, E44, F47, I60, W62, F67, Y70 | 226 |
| 2 | 1 | 13 | I11, L20, F61, I108, L284, L290, L299 | 138 |
| | 2 | 5 | I108, A109, L284, L285, L290, V293, L299, V302 | 118 |
| | 3 | 10 | I11, K15, L20, F61, I108, L262, L284, V293, L299 | 110 |
| 3 | 3 | 2 | W129, I132, L147, L198, H203, M204, I226, P248 | 103 |
| | | 11 | W129, I132, L147, F149, L160, L198, H203, M204 | 100 |
| | | 17 | W129, I132, L147, F149, T157, L160, L198, M204 | 105 |
| 4 | 1 | 11 | T157, I161, V181, L192, V343, A346, L361 | 113 |
| | 3 | 1 | I161, V183, L192, V343, A346, V347, L361 | 127 |
| | | 13 | W158, I161, V183, A188, L192, V343, A346, L361 | 103 |

**Supplementary Table 2 | Residues comprising $O_2$-trapping subareas.** The residue set with the longest $O_2$-trapping time for each subarea was used for the DFT calculations in Fig. 5.

#### Supplementary Videos

**Supplementary Video 1 | Representative $O_2$-trapping events.** The video showing relatively long $O_2$-trapping events were created using MD simulation snapshots obtained from VMD program (one frame per 100 ps). A smoothing window size of 2 frames was applied to all atoms, and the video speed was set to 80 frames per second. The simulation time span for the video corresponds to 620–700 ns of simulation #1. In the video, MBP is shown as a yellow cartoon representation. The oxidized residues detected from the LC MS/MS and the Trp residues which are near the trapped $O_2$ trajectories are shown as van der Waals surface representation and colored red and blue, respectively. $O_2$ is also shown as van der Waals surface representation and colored black. For clear visualization of the $O_2$-trapping events, water molecules, chloride and potassium ions, and other $O_2$ molecules except for the trapped $O_2$, are not shown.

**Supplementary Video 2 | Representative maltose-binding events.** The video was created using MD simulation snapshots obtained from VMD program (one frame per 1 ns). A smoothing window size of 2 frames was applied to all atoms, and the video speed was set to 25 frames per second. The simulation time span for the video corresponds to 2950–3150 ns of the simulation. In the video, MBP is shown as a yellow cartoon representation. The maltose-binding site is shown as transparent surface representation and colored blue. Maltose is shown as van der Waals surface representation and colored red. Chloride and potassium ions are shown as ball-and-stick representation. Water molecules are not shown for clear visualization of the maltose-binding events.
